# Differences in the activities of domain-swapped chimeras of two homologous GH57 glucanotransferases suggest that a glucan-binding DUF could influence donor substrate specificity

**DOI:** 10.1101/2023.08.25.554918

**Authors:** Arpita Sarkar, Pallavi Kaila, Purnananda Guptasarma

## Abstract

Five enzymes of the archaeal hyperthermophilic family of disproportionating GH57 4-α-glucanotransferases have been studied till date. Our focus here lies upon three homologous members of this family: (i) PfuAmyGT from *Pyrococcus furiosus* (PF0272), (ii) TonAmyGT from *Thermococcus onnurenius* (B6YUX8), and (iii) TliAmyGT (TLGT) from *Thermococcus litoralis* (O32462). The polypeptide chain of each of these enzymes is approximately 655 residues long, folded into three distinct domains (1, 2 and 3), and assembled into a homodimer. Domain 1 is a beta/alpha barrel containing an aspartate known to function as a catalytic nucleophile in TLGT. Domain 2 (which is helical) and domain 3 (made up of beta sheets) are thought to be domains of unknown function (or DUFs). In PfuAmyGT and TonAmyGT, we have recently identified a catalytically-important aspartate upon a loop in domain 2. In PfuAmyGT, we demonstrate the presence of two additional catalytically-important (glutamate) residues in domain 1, in a companion paper. In this paper, our focus lies upon domain 3 which hosts a second binding site (SBS) for a glucan, at its domain-domain interface with domain 2. Using strategies involving studies of both (a) domains (or pairs of contiguous domains) extracted from PfuAmyGT, and (b) chimeric three-domain enzymes recombining analogous domains between PfuAmyGT and TonAmyGT, we demonstrate that domain 3 determines the choice of the preferred glucan that acts as a donor in the glucan transfer reaction.

## Introduction

The GH57 family of enzymes is a family of glycosyl hydrolases found mainly in thermophile bacteria and hyperthermophile archaea.^1–5^ Members of this family include (a) α-amylases,^6–8^ (b) amylopullulanases,^9–11^ (c) branching enzymes,^1,12–14^ (d) 4-α-glucanotransferases,^15–21^ and (e) α-galactosidases.^22^ The polypeptide chains of certain hyperthermophilic archaeal 4-α-glucanotransferases of this family tend to be ∼665 amino acids in length,^3,20,21^ and folded into three distinct structural domains. Chains are assembled into ‘head-to-tail’ homodimers. Within these homodimers, subunits interact primarily through domain 1.^3^ An archetypal GH57 hyperthermophilic archaeal 4-α-glucanotransferase is TLGT from *Thermococcus litoralis*.^3^ Structures for TLGT (PDB IDs: 1K1X/1K1Y/1K1W) have been determined both with, and without, any bound glucans. Supplementary Fig. S1 shows a ribbon diagram representation of the TLGT (1K1X) homodimer. From this figure, it becomes evident that the TLGT dimer is primarily stabilized by inter-subunit contacts involving domain 1 from each subunit.

TLGT is homologous to 4-α-glucanotransferases from *Pyrococcus furiosus*, and *Thermococcus onnurineus,* two other archaeal hyperthermophiles. We refer to these enzymes as PfuAmyGT,^20^ and TonAmyGT,^21^ because both enzymes display the ability to function upon starch [displaying an amylase (or Amy) function], as well as upon any single (pure) malto-oligosaccharide [displaying dis-proportionating glucanotransferase (or GT) function]. With either substrate, both PfuAmyGT and TonAmyGT generate entirely similar products in the form of pools of glucose and small malto-oligosaccharides ranging in length from maltose to maltohexaose, with some small amounts of maltoheptaose and occasionally even larger species also present and detectable. The oligosaccharides in such pools differ in length amongst themselves by a single glucose unit, suggesting that the transferred sugar is glucose itself, rather than maltose or something longer.

Aligned amino acid sequences of TLGT, PfuAmyGT and TonAmyGT are shown in Fig. 1A. The percentage similarities between their sequences are shown in Fig. 1B. From both figures, it would appears that TLGT, PfuAmyGT and TonAmyGT are so homologous that they are essentially organismal sequence-variants of the same glucanotransferase. Therefore, we refer presumptively to TLGT as TliAmyGT in rest of the text. Supplementary Figs. S2 A, B and C show that PfuAmyGT and TonAmyGT have highly homologous sequences for domains 2 and 1, with sequence differences seen mainly in domain 3. In Fig. 1A, domain 1 is seen to be a little over than ∼310 amino acids long, and folded into a beta/alpha (TIM) barrel. Domain 1 is thought to be primarily responsible for activity in TliAmyGT, since it hosts a catalytic nucleophile in the form of a glutamate residue (E123) that has been demonstrated to be involved in glucan transfer.^23^ This catalytic nucleophile is conserved in the two other enzyme homologs [E131 in PfuAmyGT; E121 in TonAmyGT]. Domain 2 is ∼80 amino acids long, and made up entirely made up of alpha helical structure, while domain 3 is ∼250 amino acids long, and made up entirely of beta sheet structure. These two domains are annotated as domains of unknown function (DUFs).^24^ Supplementary Fig. S1 shows that the folded forms of domains 1, 2 and 3, are arranged in a linear fashion in each subunit of TliAmyGT, with domains 1 and 3 sandwiching domain 2, and with few inter-subunit contacts between domains 1 and 3, and no intra-subunit contacts between these two domains. We are intrigued by the presence of the DUF domains 2, and 3, in these enzymes and would like to gain insights into their functions.

**Fig. 1.**
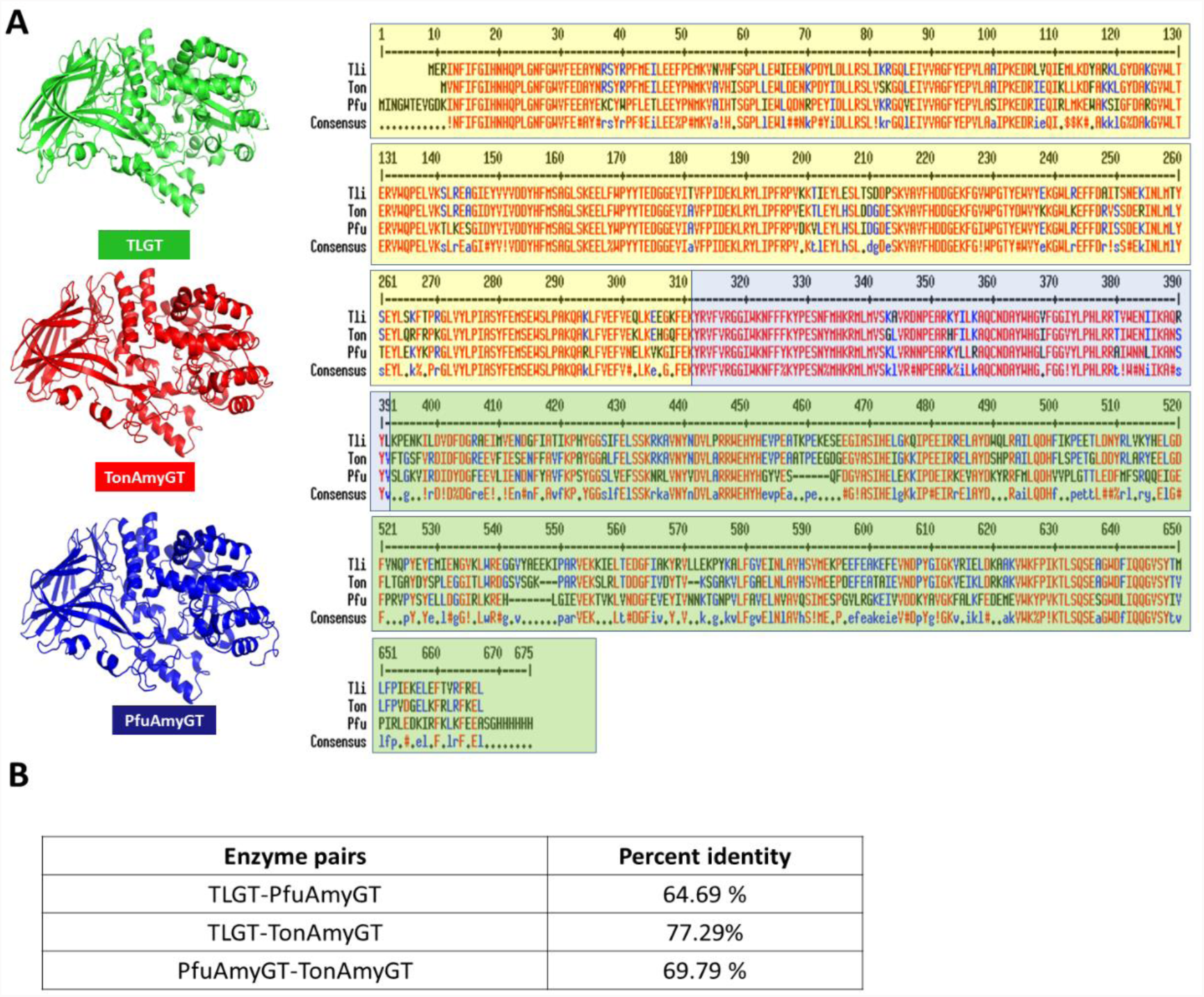
**A.** The aligned sequences of TLGT, PfuAmyGT and TonAmyGT, with domain boundaries marked. **B.** Percent identities amongst the three enzymes.

The known structure of TliAmyGT, and the modelled structures of PfuAmyGT and TonAmyGT [using i-Tasser software,^25^ and obtained through homology-based modelling using TliAmyGT as the template] are shown in Fig. 1. Our group has recently studied PfuAmyGT and TonAmyGT and found some evidence of the importance of domains 2 and 3 in the functioning of these enzymes. <u>For domain 2</u>: In both PfuAmyGT,^20^ and TonAmyGT (unpublished observations), we have found that domain 2 contains a loop which hosts a second catalytically-important aspartate residue [D362 in PfuAmyGT; D352 in TonAmyGT]. Replacement of D362 by alanine in PfuAmyGT,^20^ as well as replacement of D352 by alanine in TonAmyGT, destroys glucanotransferase activity without any detectable changes in secondary, tertiary or quaternary structure.^20^ <u>For domain 3</u>: We have also previously pointed out,^20^ that the published structure (1K1Y) of TliAmyGT reveals that a maltose of unknown origin is bound to domain 3,^3^ in the vicinity of the interface between domains 2 and 3. The presence of this maltose was left unexplained by the authors who published the enzyme’s structure; however, we have proposed that the maltose derives from the cytoplasm of the *Escherichia coli* expression host cell, and that it binds to an additional glucan-binding site in the enzyme which is located upon domain 3, i.e., over and above any glucan binding occurring in domain 1. It may be noted that such surface (second) binding sites (SBSs) for glucans are now being reported in numerous glycosidases.^2,26–30^

Given the above, our interest in this paper lies primarily in obtaining further insights into the functioning of domains 2 and 3 in GH57 enzymes from hyperthermophile archaea. The approach we have used here is to (i) produce the domains of a single GH57 glucanotransferase like PfuAmyGT, as isolated individual domains (Pfu1, Pfu2 and Pfu3), or as pairs of genetically fused natural-neighbour domains (Pfu1-Pfu2 or Pfu2-Pfu3), and also (ii) produce the domains of two different GH57 glucanotransferases (PfuAmyGT and TonAmyGT) as fusions of the three domains, 1, 2 and 3, using both the natural and wild-type order of domains (Pfu1-Pfu2-Pfu3 or Ton1-Ton2-Ton3) as well as six unnatural chimeras of domains sourced from both enzymes (e.g., Pfu1-Ton2-Pfu3 or Ton1-Pfu2-Ton3 or Pfu1-Ton2-Ton3 and so on). Our objective in making such engineered protein constructs was to examine how the enzyme’s amylase and (dis-proportionating) glucanotransferase functions involving residues E131 (in PfuAmyGT) or, potentially also E121 (in TonAmyGT) in domain 1, and D362 (in PfuAmyGT) or D352 (in TonAmyGT) in domain 2, tend to be influenced by domain 3. Given the very high level of conservation of sequence in domain 2 (∼80 % between any two enzymes), our analyses of the micro-structural features of the interface between domains 2 and 3 suggest that chimeras recombining domains from PfuAmyGT and TonAmyGT might fold and function.

Fig. 2A presents a view of a single subunit in the modelled structure of PfuAmyGT, in which the subunit is lined up in such a manner that domain 1 faces the reader and domain 3 retreats into the page. In this view, the surface of domain 3 is eclipsed by the ribbon diagram representations of domains 1 and 2. Using structural superimpositions to obtain similar views of PfuAmyGT and TonAmyGT, we show views of domain 3 in the three enzymes, now uneclipsed by domains 1 and 2, in Figs. 2B, C and D, respectively, for PfuAmyGT, TonAmyGT and TliAmyGT. From these views, it is evident that the micro-structural features of the surface of domain 3, involving its interface with domain 2, are largely conserved. This augurs well, e.g., for the potential assembly of domain 2 of PfuAmyGT with domain 3 of TonAmyGT, or e.g., for the assembly of domain 2 of TonAmyGT with domain 3 of PfuAmyGT, and so on, in chimeric (protein-engineered) enzymes that recombine domains between PfuAmyGT and TonAmyGT.

**Fig. 2.**
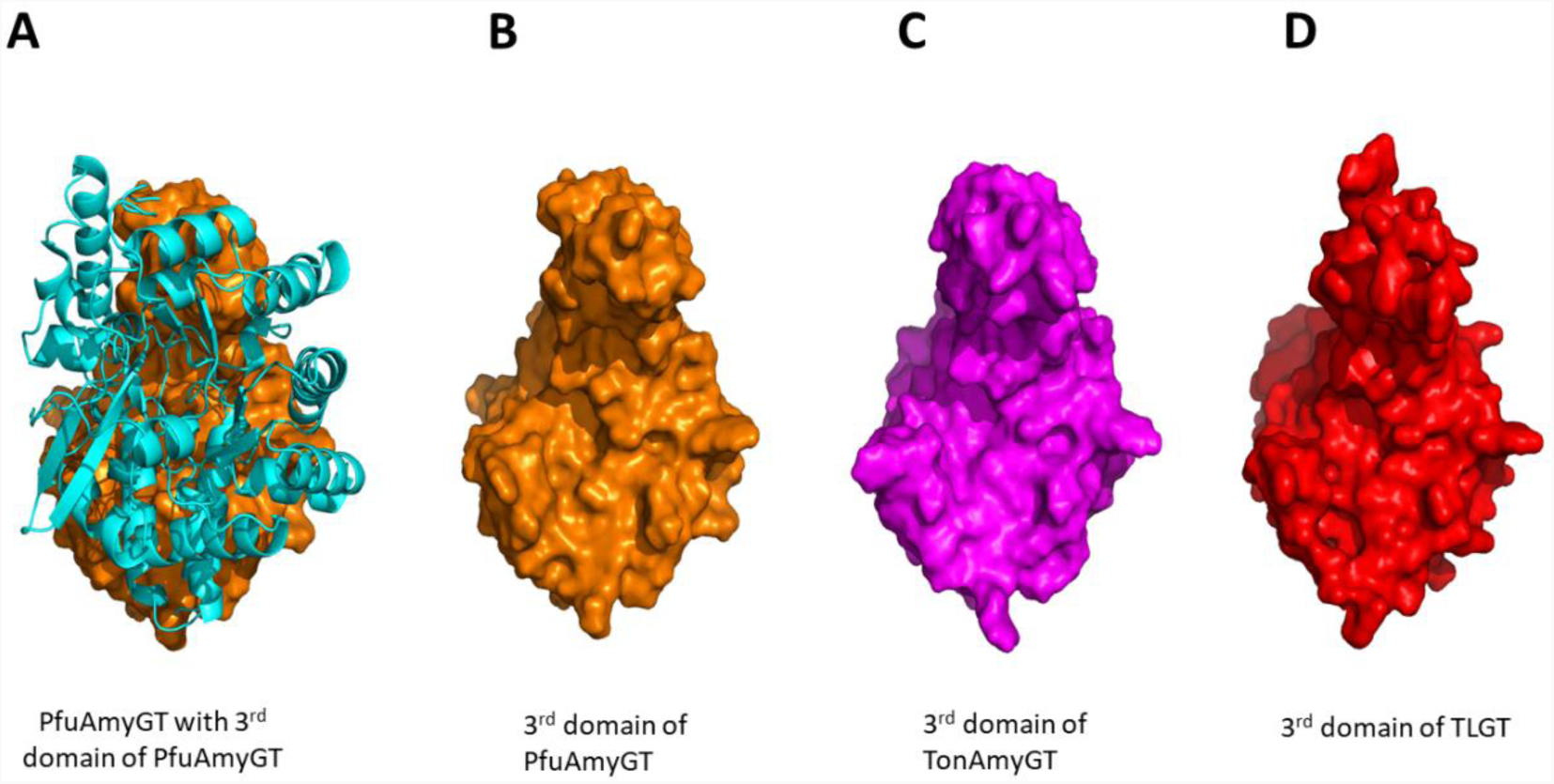
**A.** A superimposed view of domain 3 of PfuAmyGT (in a surface view representation) and domains 1 and 2 of PfuAmyGT (in a ribbon-diagram representation). **B, C and D** shows the surface of domain 3 in PfuAmyGT, TonAmyGT and TLGT, uneclipsed by ribbon-diagram representations of domain 1 and 2.

In the results shown in this paper, we ask whether domain 3 plays any role in respect of the use of the donor malto-oligosaccharide by any GH57 enzyme. We do this both by studying whether such an enzyme can function without domain 3, and by using enzymes incorporating domain 3 from a different homologous enzyme, with the hope of examining whether domain 3 somehow determines substrate specificity. Below, we show that domain 3 is indeed critical for catalytic activity in PfuAmyGT, and that it determines substrate specificity in chimeras of PfuAmyGT and TonAmyGT.

## Materials and Methods

### Bioinformatic tools

Pfam online server was used to determine the function of the three domains of PfuAmyGT (Uniprot locus ID PF0272).^24^ For sequence alignment, the Multialign online server was used.^31^ The PyMOL Molecular Graphics System, Version 1.2r3pre, Schrödinger, LLC, n.d.,^32^ was used for visualizing protein structures and for alignment of the structures. I-Tasser,^25^ and Swiss model,^33^ web services were used to model the structure(s) of PfuAmyGT, TonAmyGT and the chimera.

### Materials

Starch and glucose were procured from HiMedia, India. Malto-oligosaccharide standards of different lengths were procured from Merck (Sigma) USA. Isopropyl-1-thio-D-galactopyranoside (IPTG) was purchased from HiMedia or BR Biochem Life Sciences Pvt. Ltd., India. Restriction enzymes of the Fast Digest variety were procured from Thermo Fisher Scientific. DNA polymerase and DNA ligase were purchased from New England Biolabs, U.S.A. Silica gel 60 F254 thin layer chromatography (TLC) plates were procured from Merck, USA. Kits for plasmid isolation, DNA extraction from gels, and PCR product clean-up were obtained from Qiagen, Germany. Lauroyl sarcosine, sodium salt was obtained from Hi-Media. Guandine Hydrocholride was obtained from either Hi-Media, Promega or Merck (Sigma).

### Cloning and expression of PfuAmyGT, TonAmyGT and their domain-swapped chimeras

The gene encoding PfuAmyGT (Uniprot locus ID PF0272) was re-amplified from a synthesized gene already optimized for expression in *E. coli* and available with us.^20^ The gene was cloned in the pET-23a vector between NdeI and XhoI restriction sites. The gene encoding TonAmyGT gene (Uniprot locus ID B6YUX8) was re-amplified from a gene already available with us which was amplified from the genomic DNA of *Thermococcus onnurenius* (Kaila et al, 2019b). The gene was cloned in the pET-23a vector between NheI and XhoI restriction sites. The vector was transformed into *E. coli* XL-1 Blue cells, re-isolated and transformed into *E. coli* BL21 Star (DE3) pLysS or Rosetta (DE3) cells for expression. The domain-swapped chimeras of PfuAmyGT and TonAmyGT are described in Supplementary Tables ST1A and ST1B, which were generated using SOE-PCR (using DNA primers with sequences described in Supplementary Information Table ST2).

### Determination of protein concentration

The amino acid sequences of each of the eight proteins used (PfuAmyGT, TonAmy, and six chimeras) are known. Thus, their aromatic amino acid contents are known in terms of the numbers of the three aromatic amino acids (tryptophan, tyrosine and phenylalanine) present in their sequence. This information was used on the Expasy Protparam web server,^34^ to determine the extinction coefficient of each protein construct, and a value for the optical density at 280 nm for protein concentrations of 1 mg/ml

### Heat Treatment and affinity-based purification of PfuAmyGT, TonAmyGT and their domain-swapped chimeras

Transformed BL21 Star (DE3) pLysS or Rosetta (DE3) cells expressing PfuAmyGT, or TonAmyGT, and their domain-swapped chimeras were used to set up primary cultures containing the appropriate antibiotics (100 µg/ml ampicillin and 35 µg/ml chloramphenicol) with shaking at 220 rpm, for 12-16 hours, at 37 °C. This culture was used to inoculate a larger volume (1 L) of LB broth containing the same antibiotics and was allowed to grow until an O.D_600_ of 0.6, at which point IPTG was added to a final concentration of 1 mM to induce expression of the enzymes and their chimeras. Cultures were grown for a further for 6-7 hours at 37 °C, or overnight at 25 °C, cells were harvested through centrifugation at 7025 *g*, re-suspended in non-denaturing lysis buffer (Qiagen) containing 10 mM Tris, 50 mM NaH_2_PO_4_ and 300 mM NaCl, with a pH of 8.0, and lysed by sonication. The harvested cells of TonAmyGT and Chimera 4 were re-sususpended with non-denaturing lysis buffer (Qiagen) and 0.2 % of Sarcosyl and kept overnight for re-suspension. The lysate for the wild type enzymes and the chimeras were heated at 90 °C for 15-20 min, to thermally-aggregate and precipitate the bulk of cytoplasm-derived proteins which were further removed, along with cell debris, through centrifugation at 11290 *g* for 1 h. The supernatant, enriched in the hyperthermophile enzymes (PfuAmyGT, Chimera 1, 2, 3, 5 and 6) were then passed through Ni-NTA resin, and IMAC affinity chromatography was performed according to standard protocols (Qiagen). The supernatant of TonAmyGT and Chimera 4 were flash dialyzed against native lysis buffer and then subjected to Ni-NTA IMAC affinity chromatography. The elution from IMAC affinity chromatography was subjected to Size Exclusion Chromatography (SEC) for further purification. Eluted fractions were run on SDS PAGE gel and fractions having the enzymes were pooled together and used for further experiments. Supplementary Fig. S3 show the gel image of the pooled fraction for each enzyme.

### Determination of quaternary structural content

The formation of quaternary structure in the purified enzymes was determined through size exclusion chromatography (SEC), which was carried out on an AKTA Purifier-10 chromatographic system (GE Healthcare). For this, a was equilibrated with 50 mM Tris, pH 8.0 buffer, using a flow rate of 0.5 ml/min, and 0.2 mg/ml of each enzyme was injected and the respective chromatograms were plotted and analysed.

### Determination of secondary structural content

Circular Dichroism (CD) spectroscopy was performed on either MOS-500 (Biologic, France) or Chirascan (Applied Photophysics, UK) spectropolarimeters fitted with a Peltier unit for sample temperature control. A quartz cuvette of 0.1 cm path-length was used for collecting far UV CD spectra (200 to 250 nm) of the enzymes and their chimeras (0.2 mg/ml of each enzyme). Mean residual ellipticity (MRE) was calculated from raw ellipticity measured in millidegrees using the equation, [Ѳ] = {Ѳ (observed in mdeg) X 100 X MRW}/{1000 X protein concentration (mg/ml) X path length (cm)}, where mean residue weight (MRW) was calculated by dividing the molecular weight of the enzyme (in Daltons) by the number of amino acids in its polypeptide chain.

### Confirmation of folding and presence of tertiary structure

The presence of tertiary structure in the enzymes and their chimeras was assessed by monitoring the pear emission wavelength in the intrinsic tryptophan fluorescence of the enzyme, using a Cary Eclipse fluorimeter (Varian, USA) to examine whether the emission peak was below 350 nm, which is characteristic of free tryptophan in water and suggestive of the burial of tryptophan residues away from the aqueous solvent. Fluorescence emission spectra were recorded using excitation at 295 nm and collecting emission spectra in the range of 300-500 nm. The slit widths used for experiment was 5 nm for both excitation and emission monochromators, and the scan speed used was 100 nm/min.

### Determining size of PfuAmyGT with Gu-HCl through DLS

For determination of the hydrodynamic radius of the PfuAmyGT with Guanidine Hydrochloride (Gu-HCl), 2 mg/ml of the enzyme was incubated with the appropriate amount of 100 mM and 1 M of Gu-Hcl, at 90 °C for 12 hour and then subjected to size-analysis on a Zetasizer Nano ZS90 model dynamic light scattering instrument (Malvern Instruments). All the components were first filtered through a 0.22-micron polyvinylidene difluoride syringe filter (Millipore).

### Determination of quaternary structural content

The quaternary structure of PfuAmyGT in presence of various concentration of Gu-HCl was determined through size exclusion chromatography (SEC), which was carried out on an AKTA Purifier-10 chromatographic system (GE Healthcare). For this, a Superdex-200 Increase 10/300 GL column. The column was equilibrated with 50 mM Tris and appropriate concentration of Gu-HCl pH 8.0 buffer, using a flow rate of 0.5 ml/min, and 0.2 mg/ml of each reaction (which was pre-incubated overnight at 25 °C or 90) was injected and the respective chromatograms were plotted and analysed

### Enzymatic assay of PfuAmyGT, TonAmyGT and domain swapped chimera

PfuAmyGT, TonAmyGT and the six chimeras were examined for activity by using maltose and maltotriose as sole substrate. 20 mM of maltose/maltotriose was used with 0.2, 0.5, 1 and 2 µM of the enzyme. This reaction mixture was incubated at 90 °C for 12 h and then 2.5 µl from each sample was run on TLC along with marker consisting of 20 mM of glucose, maltose, maltotriose, maltotetraose, maltopentoase and maltoheaxose.

### Examining glucose inhibition by the enzyme in presence of maltose and maltotriose

The effect of glucose was studied with 1 µM of the enzymes in the presence of maltose/maltotriose. Glucose range was from 2, 5, 10, 15, 20, 50 and 100 mM. The reaction was carried out at 90 °C for 12 and then 2.5 µl from each sample was run on TLC along with marker.

### Examining the temperature range of the enzyme

1 µM of each enzyme was incubated at different temperature ranging from 20 °C to 100 °C and 2.5 µl from each sample was run on TLC along with marker.

### Thin Layer Chromatography (TLC) assessment of enzyme activity

The presence of glucanotransferase activity of PfuAmyGT, TonAmyGT and the chimeras was determined using thin layer chromatograms (TLCs). The different products formed by the enzymes by utilizing a single substrate (maltose or maltotriose) can be separated by TLC, using a highly polar absorbent silica (Silica plates) as stationary phase and a concoction of solvents in this particular ratio i.e. butanol:ethanol:water::50:30:20. The oligosaccharides formed move on the TLC plates according to their polarity with the mobile phase and stationary phase. These products were then visualized by spraying the plates with a mixture of methanol and sulphuric acid in the ratio of 95:5 and then heated at 120 °C for 2-5 minutes.

## Results and Discussion

Below, we present results of experiments involving PfuAmyGT, or TonAmyGT, as background controls for subsequent experiments involving protein engineered forms of these enzymes. The data in the section presented immediately below is provided for comparison with data presented in subsequent sections and to enable assessments of folding or catalytic activity in these enzymes. Below, we show that the native ‘tri-domain 1-2-3 forms’ of both enzymes, i.e., Pfu1-Pfu2-Pfu3 and Ton1-Ton2-Ton3, are both natively folded and functional. In subsequent sections, we present studies with (A) individual domains 1, 2 or 3, derived from PfuAmyGT [i.e., Pfu1, Pfu2 or Pfu3]; di-domains 1-2 or 2-3 derived from PfuAmyGT [i.e., Pfu1-Pfu2 or Pfu2-Pfu3]; and chimeric tri-domain 1-2-3 forms of novel enzymes created through recombination of domains 1, 2, and 3, of PfuAmyGT and TonAmyGT [i.e., six different chimeras with, e.g., Pfu1-Ton2-Pfu3 (short for PfuAmyGT-1-TonAmyGT-2-PfuAmyGT-3) representing a chimera that incorporates domains 1 and 3 from PfuAmyGT and domain 2 from TonAmyGT].

### Native tri-domains, Pfu1-Pfu2-Pfu3, and Ton1-Ton2-Ton3, are structured and active

Supplementary Figs. S5A and S5B show the structures of PfuAmyGT (or Pfu1-Pfu2-Pfu3) and TonAmyGT (or Ton1-Ton2-Ton3). From a consideration of these figures, as well as from considerations of Fig. 1, and Supplementary Figs. S2A and S2B, displaying ribbon-diagram representations of PfuAmyGT and TonAmyGT, it is immediately evident that (a) domain 1 consists of a beta/alpha barrel dominated by helical structure, (b) domain 2 is completely helical, and (c) domain 3 consists almost entirely of beta sheet structure, with considerable similarity of chain trajectories in the two enzymes, as demonstrated by the superimpositions of their backbones, shown in Supplementary Fig. S2C. Therefore, both enzymes contain a mixture of helical and sheet structures.

#### Studies of secondary structure

Supplementary Fig. S5 presents data from circular dichroism (CD), fluorescence emission spectroscopy, and size exclusion chromatography (SEC) experiments demonstrating that both PfuAmyGT and TonAmyGT have secondary, tertiary, and quaternary structural contents. Supplementary Figs. S5C and S5D show that the CD spectra of the two enzymes have overall mean residue ellipticities (MREs) that are comparable to each other, and also much lower in overall negative MRE intensity (−7,500 to −9,500 deg cm^2^ dmol^-1^) than that anticipated for pure helical secondary structure (−38,000 to −42,000 deg cm^2^ dmol^-1^). The shapes of the CD spectra of the two enzymes are also characterized by broad negative band minima, with comparable MRE values at all wavelengths between ∼210 and ∼220 nm. The fact that this is observed, instead of either two band minima at ∼208 and ∼222 nm, with MREs of −38,000 to −42,000 deg cm^2^ dmol^-1^ (characteristic of a purely alpha helical structure) or a single band minimum at ∼216-218 nm, with and MRE between −3,000 and −8,000 deg cm^2^ dmol^-1^ (characteristic of beta sheet structure), is thus in conformity with the anticipation that the folded forms of these enzymes should contain mixed helical and sheet structure.

#### Studies of tertiary structure

Supplementary Figs. S5E and S5F, respectively, show the fluorescence emission spectra of PfuAmyGT and TonAmyGT. Both enzymes are observed to emit maximally at ∼340 nm. Unfolded enzymes that expose constituent tryptophan residues to the aqueous solvent are anticipated to emit maximally at ∼353 nm. The blue-shifting of emission maxima by ∼13 nm in both spectra thus indicates the burial of tryptophan residues from the aqueous solvent, due to the known blue shift that occurs when tryptophan is in hydrophobic environments (owing to the emission of tryptophan being sensitive to the polarity of its environment). As can be seen in Supplementary Table ST3, there are 18, and 16, tryptophan residues, respectively, in PfuAmyGT, and TonAmyGT. The blue-shifting of spectral maxima in the folded forms of these enzymes by ∼13 nm, with respect to what is expected for unfolded enzyme(s), is thus indicative of the global burial of tryptophan side chains away from the solvent. By inference, this is indicative of tertiary structural content.

#### Studies of quaternary structure

Supplementary Figs. S5G and S5H show size exclusion chromatography (SEC) data obtained through gel filtration chromatographic studies of PfuAmyGT, and TonAmyGT, respectively, upon a Superdex-200 Increase (GE) gel filtration column with a bed volume of ∼24 ml, with the calibration plot for the column shown in Supplementary Fig. S4C. Supplementary Fig. S5G shows that PfuAmyGT elutes as a single species at ∼12 ml corresponding to a dimeric form of ∼156 kDa, in conformity with the expectation that PfuAmyGT should be a dimer [as observed in the crystal structure of its homolog, TliAmyGT, which is shown in Supplementary Fig. S1]. Supplementary Fig. S5H shows that TonAmyGT elutes at ∼8 ml, ∼12 ml and ∼14 ml, indicating that it is eluted out as a mixture of monomers, dimers and oligomers, unlike PfuAmyGT which elutes from the same column at ∼12 ml (corresponding to a dimer).

#### Studies of catalytic activity

Supplementary Fig. S6A presents a representative image of a thin layer chromatogram (TLC) which demonstrates the effects of incubating PfuAmyGT with maltotriose. The TLC image demonstrates the enzyme’s ability to catalyse the transformation of a single (pure) substrate, maltotriose, into a pool of shorter and longer malto-oligosaccharides that are separated into discrete bands by thin layer chromatography. Each band’s identity is established by comparing its mobility with that of a standard oligosaccharide, run as a mixed set of oligosaccharide markers in lane 6, which contains bands for glucose (G1), maltose (G2), maltotriose (G3), maltotetraose (G4), maltopentaose (G5), and maltohexaose (G6). Lane 1 shows the behaviour of maltotriose (20 mM) alone, whereas lane 2, 3, 4 and 5, respectively, correspond to incubations of 0.2, 0.5, 1.0 and 2.0 µM PfuAmyGT with maltotriose (20 mM). Through comparisons of mobility between sample bands and bands corresponding to standards/markers, it is evident that lanes 2, 3, 4 and 5 demonstrate a transformation of maltotriose into a pool of shorter and longer malto-oligosaccharides through dis-proportionating glucan transfer performed by PfuAmyGT. This TLC is thus presented to establish the existence of catalytic activity in intact (full-length) PfuAmyGT which contains domains 1, 2 and 3, i.e., Pfu1-Pfu2-Pfu3. Similarly, Supplementary Fig. S6E shows a representative TLC demonstrating catalytic activity in intact (full-length) TonAmyGT which contains domains 1, 2 and 3, i.e., Ton1-Ton2-Ton3.

In summary, we have established above that tri-domain polypeptides consisting of fused domains 1-2-3 are folded at the levels of formation of secondary, tertiary and quaternary structures, and also catalytically active as glucanotransferases. Supplementary Fig. S7 further establishes that both full-length PfuAmyGT and TonAmyGT also happen to produce pools of oligosaccharides through their action upon starch (using an amylase activity which is presumed to be coupled with glucanotransferase activity). These pools of oligosachharides are entirely similar to the pools generated through the disproportionating glucanotransferase activity which uses a single maltooligosachharide, such as maltotriose, or maltose. This combined data and the techniques used to collect it together comprise the baseline(s) or control(s) against data with other domain constructs presented below will be compared.

### Domain/di-domain constructs derived through truncation of Pfu1-Pfu2-Pfu3 also mostly adopt native-like structure(s), but display no catalytic activity

Fig. 3 presents our efforts to produce soluble and folded polypeptides corresponding to various domains and di-domains (i.e., fusions of neighbouring domains) of PfuAmyGT, in *E. coli*. The figure also presents results of assays investigating the presence of hydrolytic or glucan-transferring activity in such isolated domains or di-domains.

**Fig. 3.**
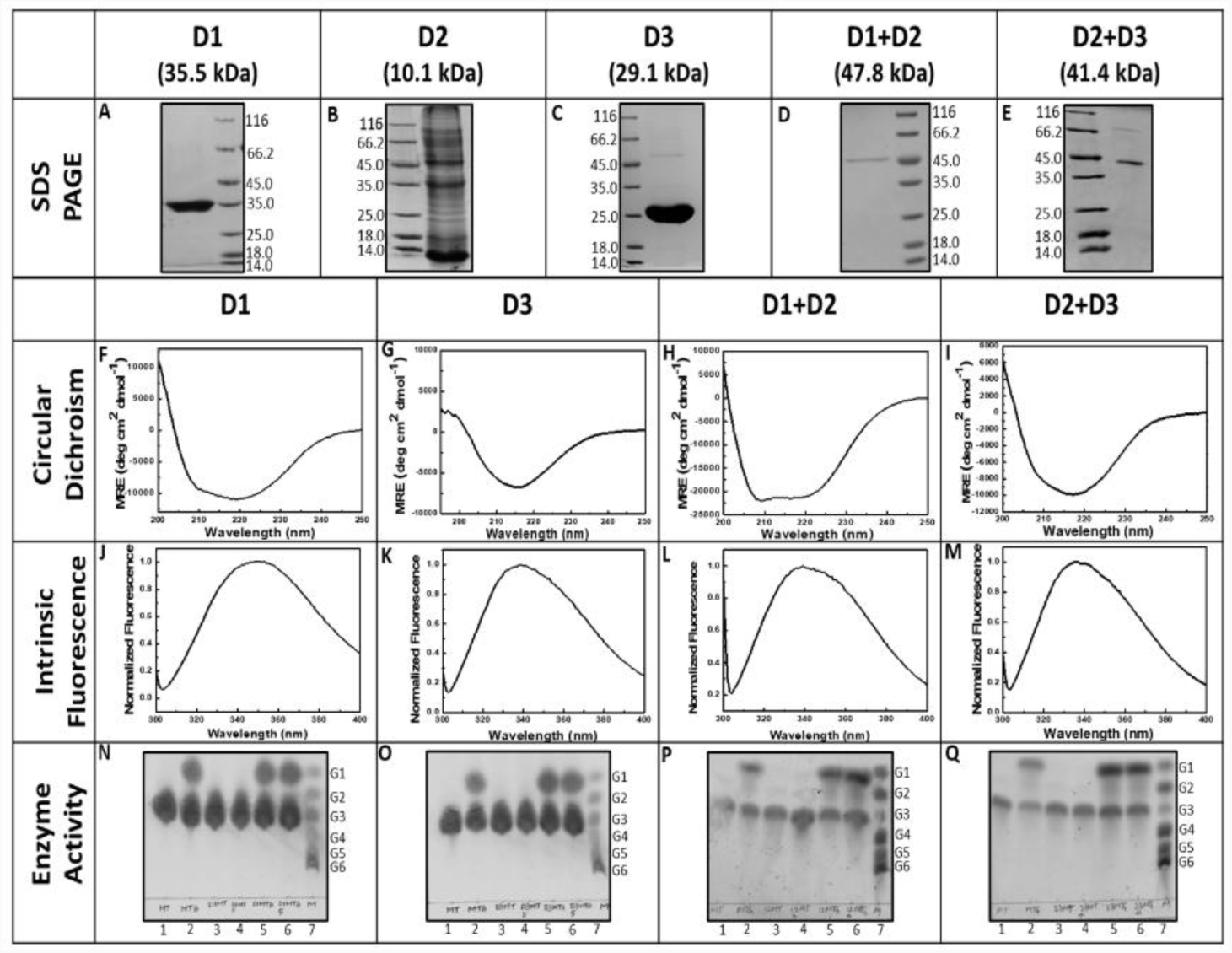
Comparative analyses of electrophoretic, spectroscopic and activity for truncated forms of PfuAmyGT consisting of Domains 1, 2 and 3, and Domain fusions 1-2 and 2-3. **A-E** SDS-PAGE profiles of IMAC-purified truncated forms**, F-I** Circular Dichroism (CD) spectra of truncated forms**, J-M** Normalized fluorescence emission spectra of truncated forms**, N-Q** Activity profiles of truncated forms. TLC data show reactions products formed through 12 h incubations of domains or domain-fusions with maltotriose at 90 °C. *Lane1*: maltotriose with no enzyme; *Lane 2*: maltotriose and glucose with no enzyme; *Lane 3*: maltotriose with 1 μMdomain/domain-fusion; *Lane 4*: maltotriose with 5 μM domain/domain-fusion; *Lane 5*: maltotriose and glucosewith 1 μM domain/domain-fusion; *Lane 6:* maltotriose and glucose with 5 μM domain/domain-fusion; *Lane 7*: premixed oligosaccharide markers containing 20 mM each of glucose (G1), maltose (G2), maltotriose (G3),maltotetraose (G4), maltopentaose (G5) and maltohexaose (G6).

#### Domains 1 and 3 (but not domain 2) are produced as soluble and folded stand-alone entities

Figs. 3A, and 3C, respectively, show SDS-PAGE images demonstrating that domains 1 (∼35.5 kDa) and 3 (∼29.1 kDa), i.e., Pfu1 and Pfu3 were overexpressed by us in amounts that were significant enough to be purified through non-denaturing affinity purification, which allowed these to be observed as pure protein gel bands of the expected size and electrophoretic mobility. In contrast, domain 2, i.e., Pfu2 could not be purified through non-denaturing affinity purification, because the overexpressed form of this domain was not soluble in the cytoplasm of the *E. coli* host. Fig. 3B thus shows instead a gel image of lysed host cell extract loaded upon a denaturing (SDS-PAGE), to show that a polypeptide of the expected size of Pfu2 (∼10.1 kDa) was overexpressed. This polypeptide could not be folded into a soluble form even after denaturing purification, ostensibly owing to a disproportionately large fraction of Pfu2’s surface being designed to remain engaged in interactions with domains Pfu1 and Pfu2 in Pfu1-Pfu2-Pfu3’s structure, since Pfu2 is natively sandwiched between Pfu1 and Pfu3, and also engaged in minor interactions with Pfu1 from the other subunit of the Pfu1-Pfu2-Pfu3 homodimer, evident from examination of Supplementary Fig. S1 (showing the homodimer of the Tli1-Tli2-Tli3 homolog, with domains highlighted in different colours). It may be surmised that domain Pfu2 has insufficient solvent-exposed hydrophilic surface for it to be soluble upon folding into native-like structure, and too many hydrophobic residues to adopt a soluble alternative structure in case folding doesn’t occur to native structure. Therefore, we think that this prevents its folding into a soluble and folded stand-alone entity. In summary, only two domains of PfuAmyGT, i.e., Pfu1 and Pfu3 were amenable to being produced, purified and examined as ‘stand-alone’ molecular entities.

#### Domains 1 and 3 appear to adopt native-like secondary structure

The polypeptide chains of both domains 1 and 3 of PfuAmyGT are characterized by CD spectra that indicate the formation of secondary structure. The type of secondary structure found to dominate the CD spectrum of these domains conformed to the type of secondary structure anticipated to be formed by each domain. Domain 1 was anticipated to form a beta-alpha barrel dominated by helices (please see Fig. 1 and Supplementary Fig. S2A). As seen in Fig. 3F, domain 1 (or Pfu1) was found to be characterized by an alpha helix-dominated CD signature displaying characteristic negative maxima at ∼208 and ∼222 nm. In contrast, Fig. 3G shows that domain 3 (or Pfu3) is characterized by a beta sheet-dominated CD signature. Once again, Fig. 1 and Supplementary Fig. S2A indicate that domain 3 is anticipated to form an all-beta structure, assuming that folding occurs to generate a stand-alone domain folded into a largely native-like structural format.

#### Domains 1 and 3 bury tryptophan residues away from the aqueous solvent to form tertiary structure

The polypeptide chains of domains 1 (or Pfu1) and 3 (or Pfu3) are characterized by fluorescence emission spectra which indicate the formation of tertiary structure. Domains 1, and 3, respectively, contain 12, and 3, tryptophan residues each, upon excitation with light of ∼280 nm (which principally excites tryptophan residues with solvent-sensitive fluorescence emission maxima), the emission spectra of domains 1, and 3, respectively, are shown in Figs. 3J, and 3K. These figures reveal that domains 1, and 3, respectively, have fluorescence emission maxima of ∼350 nm and ∼338 nm. Both emission maxima are blue-shifted with respect to ∼353 nm; the wavelength at which tryptophan residues emit maximally if they are completely exposed to an aqueous solvent. Figs. 3J and 3K thus indicate that the 12, and 3, tryptophan residues, respectively, of domains 1, and 3, happen to be substantially buried within the purified constructs, in turn indicating the folding of their polypeptide chains into collapsed tertiary structures capable of burying tryptophan residues.

#### Domain 1 appears to be a tetramer, while domain 3 is a monomer

Supplementary Figs. S4A, and S4B, show the size exclusion chromatographic (SEC) behaviour of domains 1, and 3, respectively, upon a 24 ml Superdex-200 (GE) gel-filtration column. The molecular weights of the polypeptide chains of domains 1, and 3, respectively, are 35.5, and 29.1 kDa. Although the chain size of domain 1 (or Pfu1) is not very much bigger than that of domain 3 (or Pfu3), domain 1 elutes from the Superdex-200 Increase (GE) column at a much earlier volume (∼12.2 ml) than domain 3 (∼14.6 ml). The elution volume indicates that domain 1 could be a dimer of dimers, since it appears to have a hydrodynamic volume which is substantially larger than that of a monomer, or a dimer. Indeed, as already seen in the first section, the elution volume of domain 1 (∼12.2 ml) is very close to that of the PfuAmyGT dimer (∼12 ml), even though polypeptide chain of the PfuAmyGT monomer (∼78.1 kDa) is more than twice the size of domain 1 (∼35.5 kDa), and the size of the PfuAmyGT dimer (∼156 kDa) eluting at ∼12 ml is more than four times the size of domain 1. From the structures and positions of domains in the PfuAmyGT dimer, as seen in Supplementary Fig. S1, it is clear that the subunit interface in the dimer is largely made up of contacts between the mostly helical first domains (i.e., domain 1) from each of the two subunits. Therefore, we conclude that domain 1 forms a dimer through its folding into a dimeric beta/alpha barrel structure in the same structural format in which it is present within PfuAmyGT, and that this dimer then further associates into a tetramer, i.e., a dimer of dimers. In contrast, the SEC data in Supplementary Fig. S4B suggests that domain 3 (Pfu3) is a monomer, as it displays an elution volume (∼14.6 ml) which is too large for it to be a dimer (∼13.6-14.1 ml), but small enough for it to be a monomer (expected to elute at ∼15.0-15.5 ml for a globular domain of ∼29.1 kDa). An examination of the structures and positions of domains 1, 2 and 3, in the dimeric structure of PfuAmyGT in Supplementary Fig. S2 (modelled upon TliAmyGT’s structure) indicates that no scope exists for domain 3 to form a dimer through native-like contacts, since the locations of domain 3 in the two subunits are very far apart from each other. The structure of domain 3 is also not very globular, suggesting that it could manifest a larger hydrodynamic volume than its actual volume, causing it to elute somewhat earlier. Therefore, we conclude that domain 3 is monomeric.

#### Domain 1 and domain 3 display no catalytic activity

Figs. 3N and 3O, respectively, show thin layer chromatography (TLC) data for experiments in which domains 1, or 3, were incubated for 12 h at 90 ⁰C with maltotriose. The lanes of the TLCs shown establish that domains 1 and 3 individually fail to display catalytic activity, i.e., each of these domains is incapable of transforming maltotriose into a pool of malto-oligosaccharides. In these figures, lanes 1, and 2, respectively, show the TLC behaviour of maltotriose (20 mM) alone, and maltotriose (20 mM) mixed with glucose (20 mM), in the absence of any protein domain construct, as saccharide controls. Lanes 3, and 4, respectively, correspond to incubations of 1 µM, and 5 µM, domain with maltotriose (20 mM), with these lanes in Fig. 3N corresponding to incubations of maltotriose with domain 1, and the lanes in Fig. 3O corresponding to incubations of maltotriose with domain 3, with no lane showing evidence of any glucan transferring activity. Lanes 5, and 6, respectively, correspond to incubations of 1 µM, and 5 µM, of domain 1, or domain 3, with maltotriose (20 mM) mixed with glucose (20 mM). Lane 7 shows a set of markers (G1-G6) similar to those shown in lane 6 of Supplementary Fig. S6A. It is evident from both Figs. 3N and 3O that the maltotriose incubated for 12 h at 90 ᵒC with stand-alone protein constructs comprising domains 1, and 3, respectively, does not become transformed into malto-oligosaccharide species that are either bigger, or smaller, than maltotriose, whether in the presence or absence of glucose. This firmly establishes that domains 1, and 3, are not catalytically active as glucanotransferases. Domain 1 is anticipated to contain the main catalytic centre(s) of any GH57 enzyme, while domains 2 and 3 are domains of unknown function (DUFs). Full-length PfuAmyGT contains all three domains, with domain 2 bridging domains 1 and 3.

#### Di-domains Pfu1-Pfu2 and Pfu2-Pfu3 fold to adopt secondary and tertiary structure

Figs. 3D and 3E, respectively, show that di-domains 1-2 (or Pfu1-Pfu2), and 2-3 (or Pfu2-Pfu3), were successfully overexpressed and purified as soluble stand-alone species. In di-domain 1-2, domain 2 could be expected to assemble with domain 1, leaving the construct unfulfilled in respect of contacts with domain 3. Likewise, in di-domain 2-3, domain 2 could be expected to assemble with domain 3, leaving the construct unfulfilled in respect of contacts with domain 1. Since the interfaces of domain 2 (common to both di-domain constructs) with domains 1, or 3, comprise only a small fraction of the total surface area of either di-domain construct (assuming successful folding of di-domains into native-like structural format), this explains our success in producing these two di-domains. Figs. 3H and 3I, respectively show the CD spectra of di-domains 1-2 and 2-3. Based on discussions of CD spectra already included in sections above, our comment here would be that both di-domains are well-folded. Di-domain 1-2 clearly contains significant well-folded sections of chains (with an MRE of −20,000 deg cm^2^ dmol^-1^) dominated by helical structures present in both domains. Di-domain 2-3 has a lower overall content of structure (an MRE of −7,000 deg cm^2^ dmol^-1^), a band minimum at ∼218 nm establishing that the spectrum is dominated by the beta sheet-based structure of the larger component domain (domain 3) in this fusion; however, with the small helical domain (domain 2) also establishing its presence through the visible band-shoulder at ∼208 nm, derived from its helical structure(s). Regarding the formation of tertiary structure by di-domains 1-2 and 2-3, Figs. 3L and 3M, respectively, show their fluorescence emission spectra. Once again, based on the discussions of fluorescence spectra already included in previous sections, we see that both di-domains bury away component tryptophan residues, with di-domain 1-2 displaying an emission maximum of ∼338 nm, and di-domain 2-3 displaying an emission maximum of ∼340 nm. The emission spectral envelope of di-domain 1-2 displays a shoulder at ∼350 nm. This indicates that some tryptophan residues with high fluorescence quantum yields are not fully buried in domain 2, ostensibly because of the absence of domain 3.

#### The di-domains Pfu1-Pfu2 and Pfu2-Pfu3 display no catalytic activity

Next, we shifted our attention to examining whether any catalytic activity exists in the two di-domain constructs, 1-2 (Pfu1-Pfu2), and 2-3 (Pfu2-Pfu3). Figs. 3P and 3Q, respectively, establish that di-domain constructs, 1-2, and 2-3, display no hydrolytic or glucan-transferring ability whatsoever, just like the stand-alone forms of domains 1, and 3. Our data thus suggests that domain 3 plays an accessory role, and is critical for the display of hydrolytic and glucan transferring activities by domain 1, in each of the two monomeric subunits of folded PfuAmyGT (which is a homodimer). From an examination of the structural model of the homodimeric form of the enzyme seen in Supplementary Fig. S1, it is clear that domain 3 is not a part of the subunit-subunit interface. In summary, in all of the data presented above, we have established that there is no hydrolytic or glucan-transferring catalytic activity in stand-alone forms of domain 1 (or Pfu1), domain 3 (or Pfu3), di-domain 1-2 (or Pfu1-Pfu2), or di-domain 2-3 (or Pfu2-Pfu3), although both hydrolytic and glucan-transferring activities are seen in full-length PfuAmyGT (or Pfu1-Pfu2-Pfu3). This suggests that activity is dependent upon the presence of both domains 1 and 3.

### Six different chimeric tri-domain-1-2-3 constructs that recombine domains of Pfu1-Pfu2-Pfu3 & Ton1-Ton2-Ton3 are all folded, soluble and display catalytic activity in ways that offer insights into the role of domain 3

Recombination of domains 1, 2 and 3, from the homologous GH57 hydrolases, PfuAmyGT (or Pfu1-Pfu2-Pfu3), and TonAmyGT (or Ton1-Ton2-Ton3), was attempted through the construction and expression of chimeric genetic constructs. In each of these constructs, each of the three domains was sourced from either or Pfu1-Pfu2-Pfu3 or Ton1-Ton2-Ton3. The rationale underlying the construction of such chimeras, and the likelihood of their undergoing folding as well as domain-domain assembly, and subunit-subunit assembly have already been discussed in the introductory section of this manuscript. However, as is well-known, predictions regarding the folding and function of engineered proteins can be frustrated by various factors that cannot always be anticipated. Therefore, it is necessary to produce such proteins experimentally, to examine whether they undergo folding naturally, without undergoing any aggregation or degradation, or whether they need to be refolded from chains that deposit into inclusion bodies (often with such refolding failing to occur, due to incompatibilities of side chain packing inherently caused by the engineering). We found that all of the tri-domain-1-2-3 chimeras of Pfu1-Pfu2-Pfu3 and Ton1-Ton2-Ton3 that we produced were folded and soluble stand-alone protein entities.

Recombination of three domains involving two homologous enzymes with retention of the domains in the native sequence (i.e., only as constructs using the sequence, 1-2-3, rather than any other sequence such as 3-1-2) is theoretically expected to generate a total of eight different constructs. Denoting the domain 1 of PfuAmyGT as ‘Pfu-1’, the domain 1 of TonAmyGT as ‘Ton-1’, and so on, these eight constructs consist of (A) six protein-engineered chimeras, and (B) two native protein constructs. We named and defined the six chimeras, respectively, to be <u>Chimera 1</u> (Pfu1-Pfu2-Ton3); <u>Chimera 2</u> (Pfu1-Ton2-Pfu3); <u>Chimera 3</u> (Ton1-Ton2-Pfu3); <u>Chimera 4</u> (Ton1-Pfu2-Ton3); <u>Chimera 5</u> (Pfu1-Ton2-Ton3); and <u>Chimera 6</u> (Ton1-Pfu2-Pfu3). A summary of these chimeras is provided in Supplementary Information Table ST1A and ST1B. The two native protein constructs in which domains 1, 2 and 3, occur in their natural sequence are, of course, <u>PfuAmyGT-1-2-3</u> (i.e., Pfu1-Pfu2-Pfu3); and <u>TonAmyGT-1-2-3</u> (i.e., Ton1-Ton2-Ton3), already described in the first sub-section of this section on results. Further, we produced genes encoding the remaining six (chimeric) constructs. Prior to considering the experimental results, it may be noted that the anticipated folded structures of the six chimeras were also obtained by us through structural model-building using the Swiss model website.^32^ These are shown for the different chimeras defined above, in Figs. 4A, 4B and 4C, for chimeras 1, 2 and 3, and in Figs. 5A, 5B and 5C, for chimeras 4, 5 and 6.

**Fig. 4.**
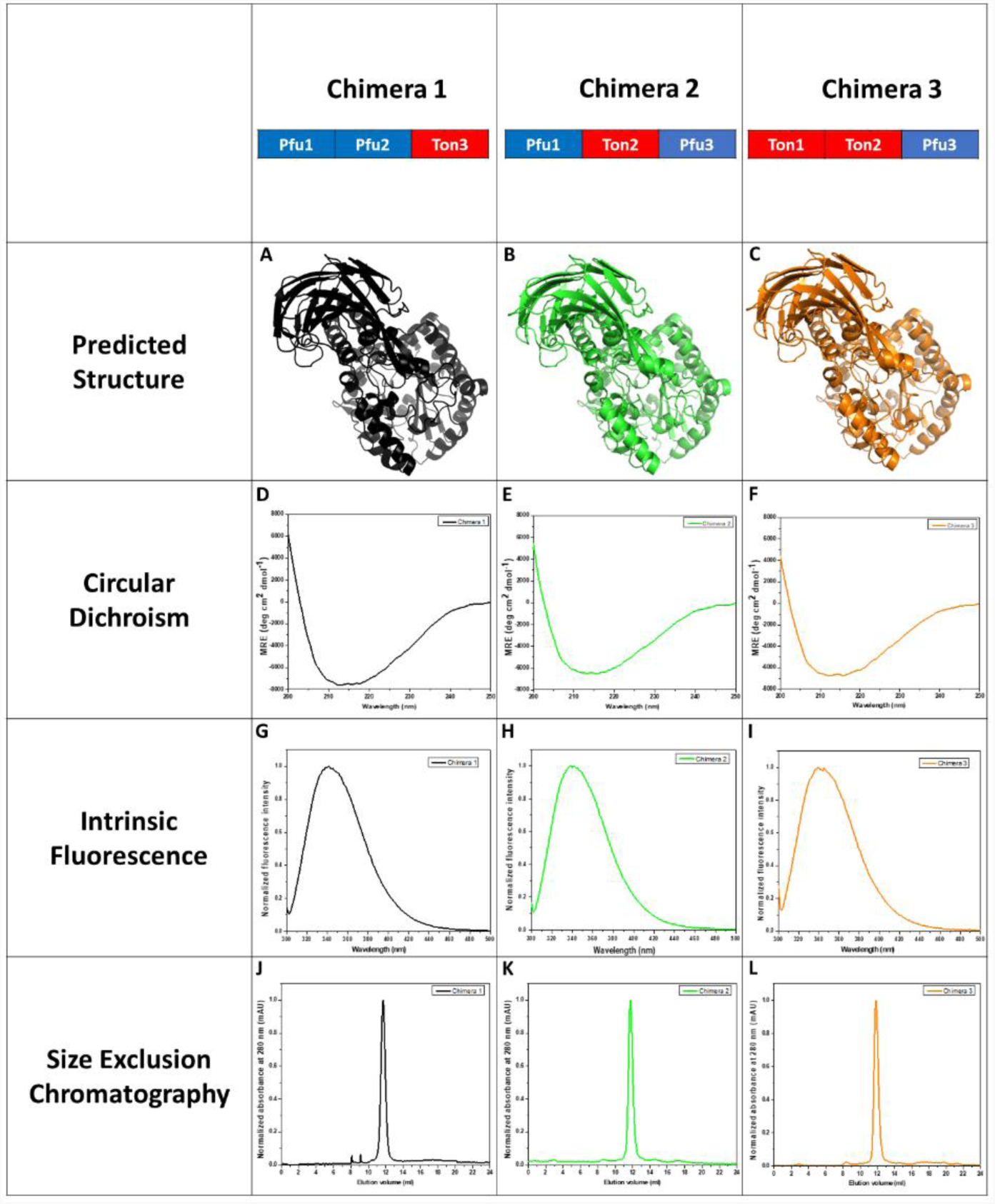
Comparative analyses of predicted structure, spectroscopic and activity for chimera 1,2 and 3. **A-C** Modelled structure of the chimeras**, D-F** Circular Dichroism (CD) spectra of the chimera**, G-I** Normalized fluorescence emission spectra of the chimeras**, J-L** Normalized gel filtration chromatogram of the chimeras.

**Fig. 5.**
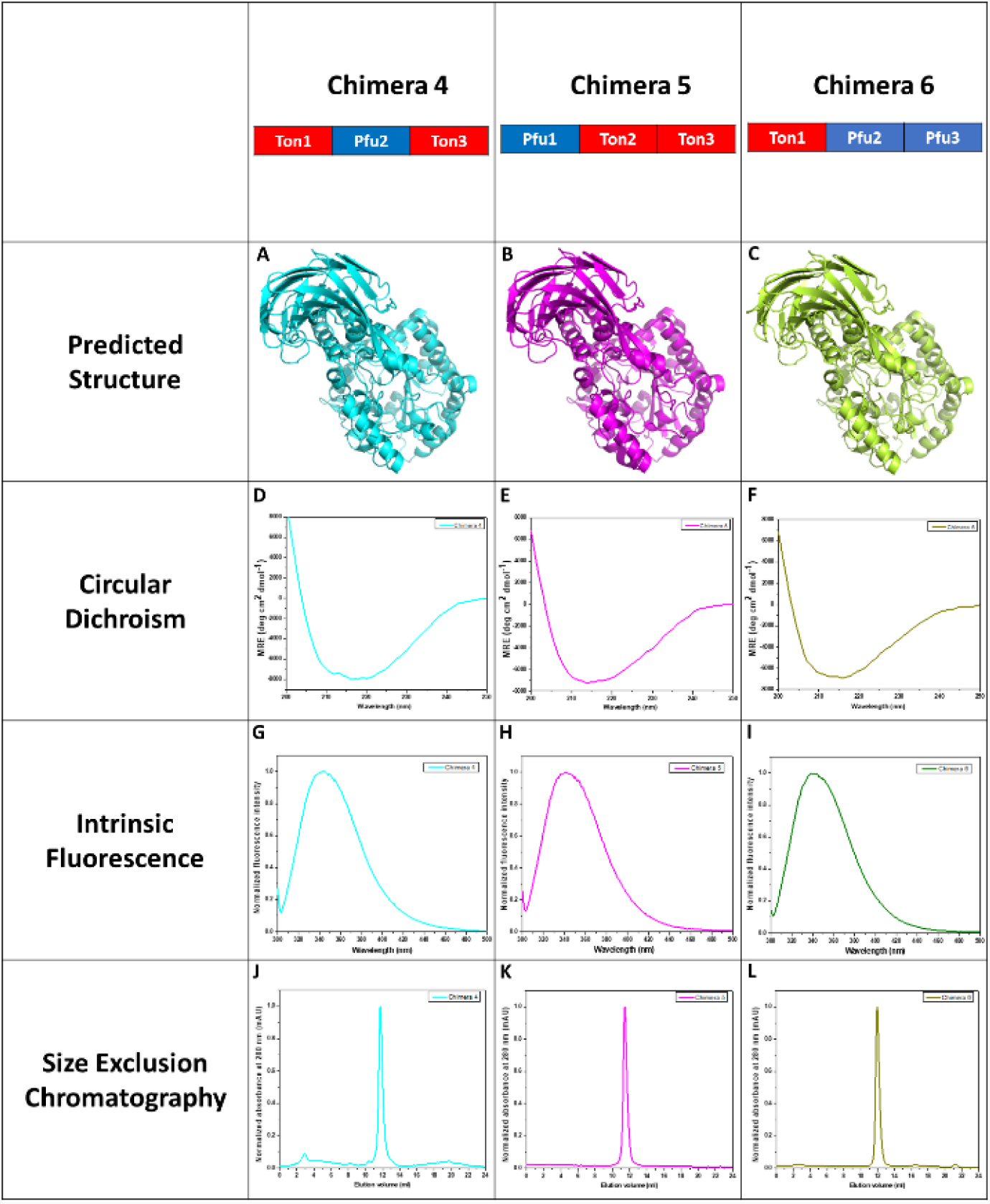
Comparative analyses of predicted structure, spectroscopic and activity for chimera 4,5 and 6. **A-C** Modelled structure of the chimeras**, D-F** Circular Dichroism (CD) spectra of the chimera**, G-I** Normalized fluorescence emission spectra of the chimeras**, J-L** Normalized gel filtration chromatogram of the chimeras.

#### The six chimeric and two native sequences of domains 1, 2 and 3 are well-folded

The CD spectra of the six chimeras are shown in Figs. 4D, 4E, 4F, 5D, 5E and 5F, respectively, for chimeras 1, 2, 3, 4, 5 and 6. We have already reviewed evidence in Supplementary Figs. S5C, and S5D, for the folding of Pfu1-Pfu2-Pfu3 and Ton1-Ton2-Ton3 comprising the remaining two constructs from the list of all 1-2-3 constructs involving both source enzymes. From the CD spectra, it is evident that each of the six chimeras folds to adopt native-like structure, in terms of the overall content of secondary structure present in each of them. Further, Figs.4G, 4H, 4I, 5G, 5H and 5I show, respectively, the fluorescence emission spectra of chimeras 1, 2, 3, 4, 5 and 6. With respect to these too, it may be noted that we have already reviewed evidence in Supplementary Figs. S5E, and S5F, for the folding of Pfu1-Pfu2-Pfu3 and Ton1-Ton2-Ton3, in terms of the fluorescence emission spectra of these constructs, and the spectral maxima displayed by their fluorescence spectra. Comparison of the fluorescence emission spectra of the six chimeras with the spectra for the two native constructs shows that they are all similar to each other. This indicates that they have all undergone folding to similar extents, and that they have buried their tryptophan residues to similar extents, since their emission maxima are blue-shifted to similar extents, with respect to the emission maximum of ∼353 nm that is seen for tryptophan residues exposed completely to the aqueous solvent. It may be noted from Supplementary Table ST3 that Pfu1-Pfu2-Pfu3 contains 18 tryptophan residues, while Ton1-Ton2-Ton3 contains 16 tryptophan residues. Therefore, the similarity of the emission spectra of all eight forms does indeed suggest that the global (molecule-wide) features of chain folding are similar in all eight cases.

#### The six chimeric and two native sequences of domains 1, 2 and 3 are also dimeric

As with the formation of secondary and tertiary structure, we found that the formation of quaternary structure too was similar for all eight constructs. Figs. 4J, 4K, 4L, 5J, 5K and 5L, respectively, show the gel filtration elution chromatograms of chimeras 1, 2, 3, 4, 5 and 6. We have already reviewed evidence in Supplementary Figs. S5G, and S5H, of the formation of quaternary structure by Pfu1-Pfu2-Pfu3 and Ton1-Ton2-Ton3, in the form of gel filtration elution or size exclusion (SEC) chromatograms. From an examination of these, it is clear that the two dimeric forms of the native constructs, Pfu1-Pfu2-Pfu3, and Ton1-Ton2-Ton3, and those of the six chimeras behave similarly during elution from a Superdex 200 Increase (GE) column of ∼24 ml bed volume. All eight constructs display an elution volume in the neighbourhood of ∼11.8-12.0 ml, corresponding to a molecular weight of ∼150 kDa. As already discussed, ∼150 kDa is close to the molecular weight of the homodimeric forms of Pfu1-Pfu2-Pfu3 and Ton1-Ton2-Ton3. In summary, the CD spectra, fluorescence spectra, and gel filtration chromatograms, respectively, establish that all six chimeras fold to adopt native-like secondary, tertiary and quaternary structure.

#### The six chimeric constructs are catalytically active in different ways, with these offering insights into the roles of domains

Qualitatively similar assays to those shown for PfuAmyGT in Supplementary Figs. S6A, B, C and D i.e., assessments of saccharide species seen in TLCs, were performed for all six chimeras. These TLCs are shown in Figs. 6 and 7. Activity assays were also performed for the two native protein constructs, Pfu1-Pfu2-Pfu3 and Ton1-Ton2-Ton3, as shown in Supplementary Fig. S6. The data in these TLCs is presented to facilitate assessment of catalytic efficiency/activity under varying experimental conditions to test glucanotransferase activity, e.g., using (i) different saccharide substrates (maltose/maltotriose), (ii) different enzyme concentrations, (iii) different temperatures, and (iv) different concentrations of glucose (to examine any inhibiting effect of glucose upon activity). Visual assessments of the raw activity data in the TLCs in Figs. 6 and 7 were then used to semi-quantitatively ‘bin’ this activity into scheme for qualitative assessment of the efficiencies of production of an equilibrium pool of saccharide species (glucose, maltose, maltotriose, maltotetraose, maltopentaose, and maltohexaose) generated through ‘dis-proportionating’ glucan transfer, beginning with either maltose, or maltotriose. The basic characteristic that was assayed was the ability of each of the eight constructs to transform the sole substrates, maltose, or maltotriose, into multiple saccharide species of different lengths, both shorter and longer than maltose, or maltotriose.

**Fig. 6.**
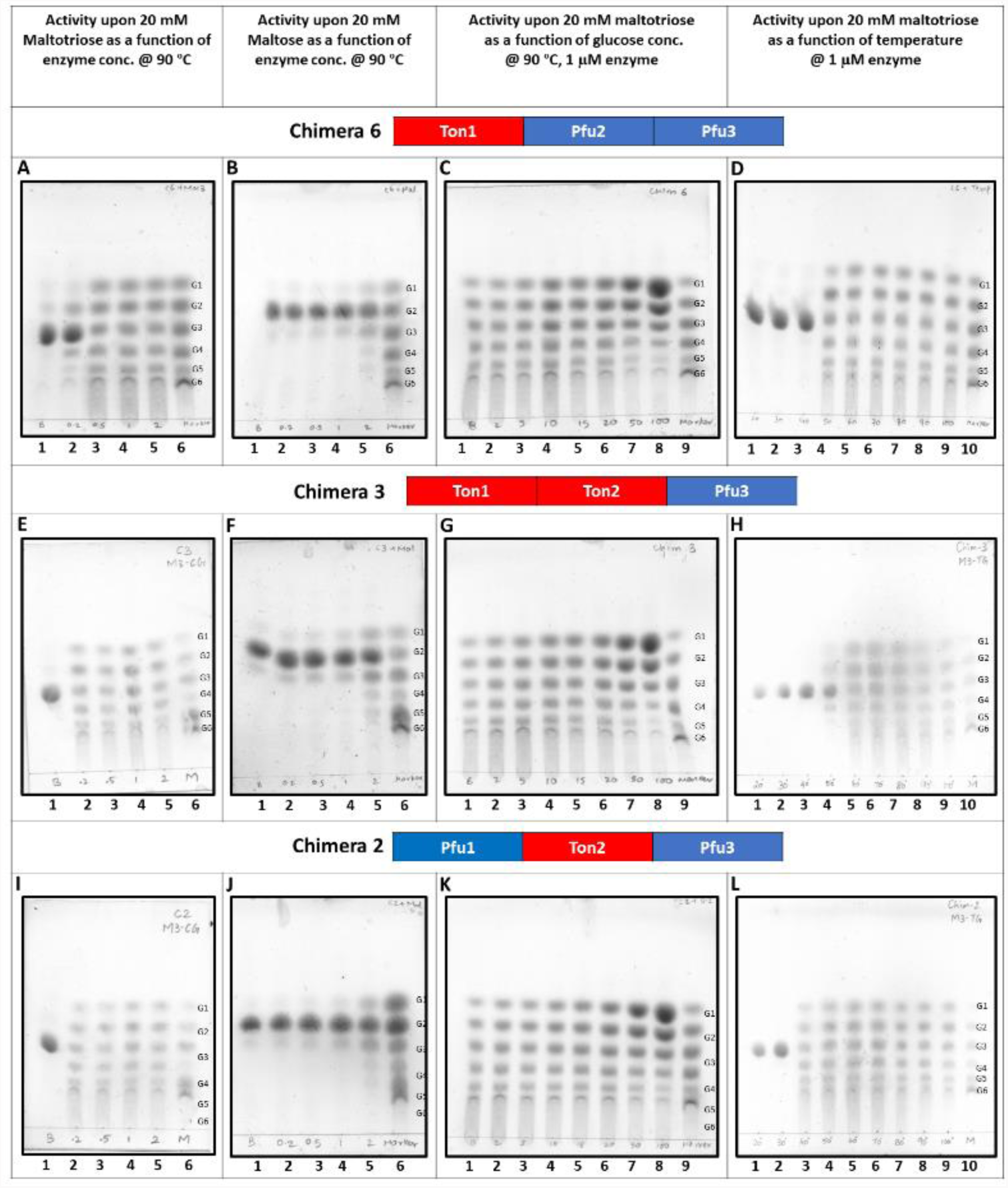
TLC experiments examining the activity profile of Chimera 6,3 and 2. **A,E and I** TLCs are showing the reaction products formed through 12 h incubations of chimera with maltotriose(20 mM) at 90 °C. *Lane 1*: maltotriose with no enzyme; *Lane 2*: maltotriose with 0.2 μM enzyme; *Lane3*: maltotriose with 0.5 μM enzyme; *Lane 4*: maltotriose with 1 μM enzyme; *Lane 5*: maltotriose with 2 μM enzyme; *Lane 6*: premixed oligosaccharide markers containing 20 mM each of glucose (G1),maltose (G2), maltotriose (G3), maltotetraose (G4), maltopentaose (G5) and maltohexaose (G6). **B, F and J** TLCs are showing the reaction products formed through 12 h incubations of chimera with maltose (20 mM) at 90 °C. *Lane 1*: maltose with no enzyme; *Lane 2*: maltose with 0.2 μM enzyme; *Lane3*: maltose with 0.5 μM enzyme; *Lane 4*: maltose with 1 μM enzyme; *Lane 5*: maltose with 2 μM enzyme; *Lane 6*: premixed oligosaccharide markers containing 20 mM each of glucose (G1),maltose (G2), maltotriose (G3), maltotetraose (G4), maltopentaose (G5) and maltohexaose (G6). **C, G and K** TLCs are showing the reaction product formed through 12 h incubations of chimera (1 µM) with maltotriose (20 mM) at 90 °C as a function of glucose concentration. *Lane 1*: maltotriose with enzyme and no glucose; *Lane 2*: maltotriose with 2 mM glucose; *Lane 3*: maltotriose with 5 mM glucose; *Lane 4*: maltotriose with 10 mM glucose; *Lane 5*: maltotriose with 15 mM glucose; *Lane 6*: maltotriose with 20 mM glucose; *Lane 7*: maltotriose with 50 mM glucose; *Lane 8* maltotriose with 100 mM glucose; ; *Lane 9*: premixed oligosaccharide markers containing 20 mM each of glucose (G1),maltose (G2), maltotriose (G3), maltotetraose (G4), maltopentaose (G5) and maltohexaose (G6). **D, H and L** TLCs are showing the reaction product formed through 12 h incubations of chimera (1 µM) with maltotriose (20 mM) as a function of temperature. *Lane 1*: maltotriose with enzyme at 20 °C; *Lane 2*: maltotriose with enzyme at 30 °C; *Lane 3*: maltotriose with enzyme at 40 °C; *Lane 4*: maltotriose with enzyme at 50 °C; *Lane 5*: maltotriose with enzyme at 60 °C; *Lane 6*: maltotriose with enzyme at 80 °C; *Lane 7*: maltotriose with enzyme at 90 °C; *Lane 8*: maltotriose with enzyme at 100 °C; *Lane 9* maltotriose with 100 mM glucose; *Lane 10*: premixed oligosaccharide markers containing 20 mM each of glucose (G1),maltose (G2), maltotriose (G3), maltotetraose (G4), maltopentaose (G5) and maltohexaose (G6).

**Fig. 7.**
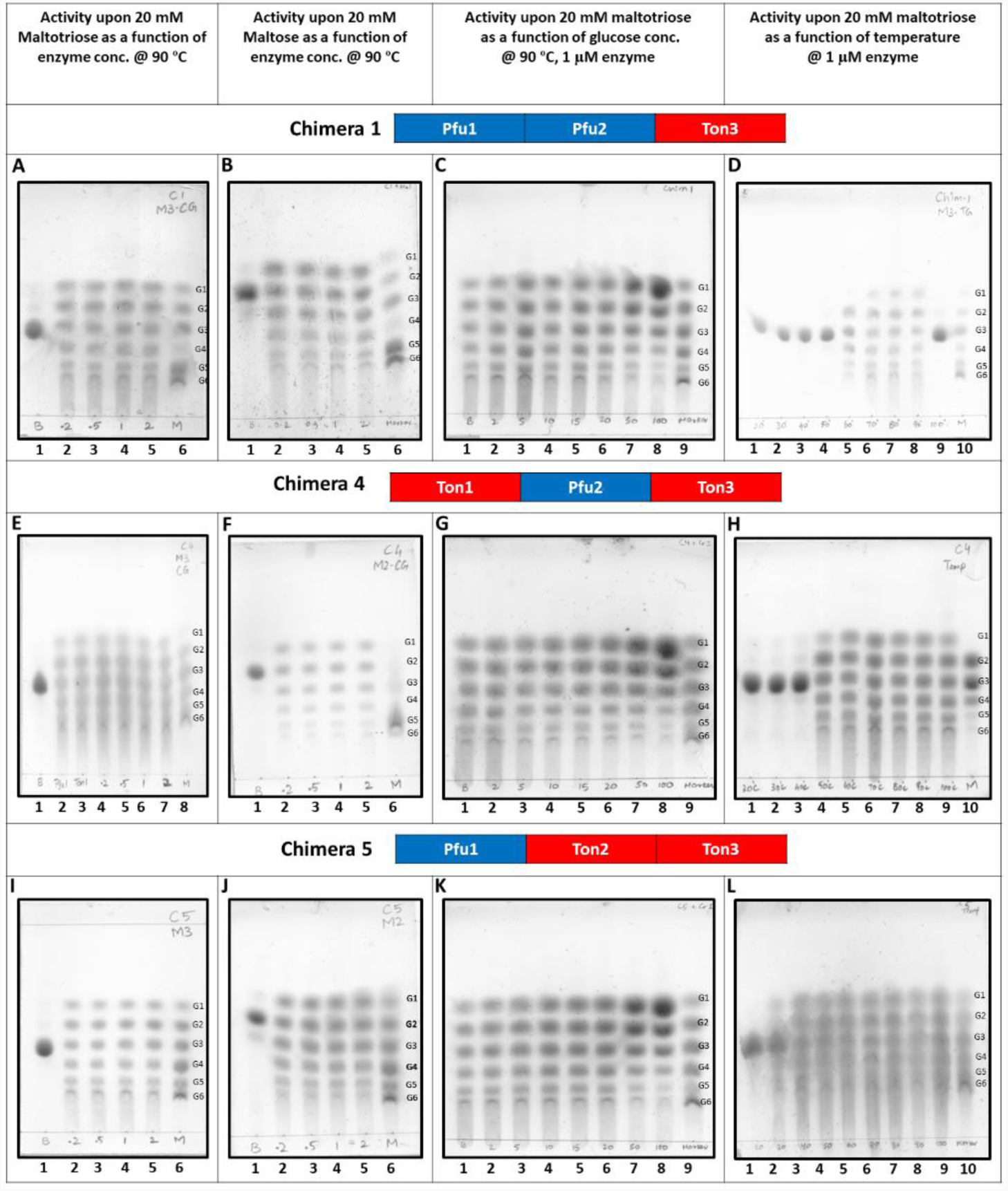
TLC experiments examining the activity profile of Chimera 1,5 and 6. **A,E and I** TLCs are showing the reaction products formed through 12 h incubations of chimera with maltotriose(20 mM) at 90 °C. *Lane 1*: maltotriose with no enzyme; *Lane 2*: maltotriose with 0.2 μM enzyme; *Lane3*: maltotriose with 0.5 μM enzyme; *Lane 4*: maltotriose with 1 μM enzyme; *Lane 5*: maltotriose with 2 μM enzyme; *Lane 6*: premixed oligosaccharide markers containing 20 mM each of glucose (G1),maltose (G2), maltotriose (G3), maltotetraose (G4), maltopentaose (G5) and maltohexaose (G6). **B, F and J** TLCs are showing the reaction products formed through 12 h incubations of chimera with maltose (20 mM) at 90 °C. *Lane 1*: maltose with no enzyme; *Lane 2*: maltose with 0.2 μM enzyme; *Lane3*: maltose with 0.5 μM enzyme; *Lane 4*: maltose with 1 μM enzyme; *Lane 5*: maltose with 2 μM enzyme; *Lane 6*: premixed oligosaccharide markers containing 20 mM each of glucose (G1),maltose (G2), maltotriose (G3), maltotetraose (G4), maltopentaose (G5) and maltohexaose (G6). **C, G and K** TLCs are showing the reaction product formed through 12 h incubations of chimera (1 µM) with maltotriose (20 mM) at 90 °C as a function of glucose concentration. *Lane 1*: maltotriose with enzyme and no glucose; *Lane 2*: maltotriose with 2 mM glucose; *Lane 3*: maltotriose with 5 mM glucose; *Lane 4*: maltotriose with 10 mM glucose; *Lane 5*: maltotriose with 15 mM glucose; *Lane 6*: maltotriose with 20 mM glucose; *Lane 7*: maltotriose with 50 mM glucose; *Lane 8* maltotriose with 100 mM glucose; ; *Lane 9*: premixed oligosaccharide markers containing 20 mM each of glucose (G1),maltose (G2), maltotriose (G3), maltotetraose (G4), maltopentaose (G5) and maltohexaose (G6). **D, H and L** TLCs are showing the reaction product formed through 12 h incubations of chimera (1 µM) with maltotriose (20 mM) as a function of temperature. *Lane 1*: maltotriose with enzyme at 20 °C; *Lane 2*: maltotriose with enzyme at 30 °C; *Lane 3*: maltotriose with enzyme at 40 °C; *Lane 4*: maltotriose with enzyme at 50 °C; *Lane 5*: maltotriose with enzyme at 60 °C; *Lane 6*: maltotriose with enzyme at 80 °C; *Lane 7*: maltotriose with enzyme at 90 °C; *Lane 8*: maltotriose with enzyme at 100 °C; *Lane 9* maltotriose with 100 mM glucose; *Lane 10*: premixed oligosaccharide markers containing 20 mM each of glucose (G1),maltose (G2), maltotriose (G3), maltotetraose (G4), maltopentaose (G5) and maltohexaose (G6).

The ‘binned’ summary of data from Figs. 6 and 7 is shown in Fig. 8, as well as in Supplementary Fig. S6. In both Figs. 8 and 9, boxes coloured ‘green’ represent a construct’s ability to engage in reactions that produce equilibrium concentrations of products characterized by equal amounts of glucose, maltose maltotriose, maltotetraose, maltopentaose and maltohexaose; boxes coloured ‘amber’ represent a construct’s ability to produce a few (but not all) of the above saccharide species over the 12 h duration of enzymatic assays conducted at 90 ᵒC; and boxes coloured ‘red’ represent a lack of any detectable activity, i.e., for instances in which there was no detectable transformation of pure maltose, or pure maltotriose, into the remaining five saccharide species. Our interpretation of this complex grid of data, and the broad conclusions drawn therefrom regarding the native and chimeric constructs are presented below.

**Fig. 8.**
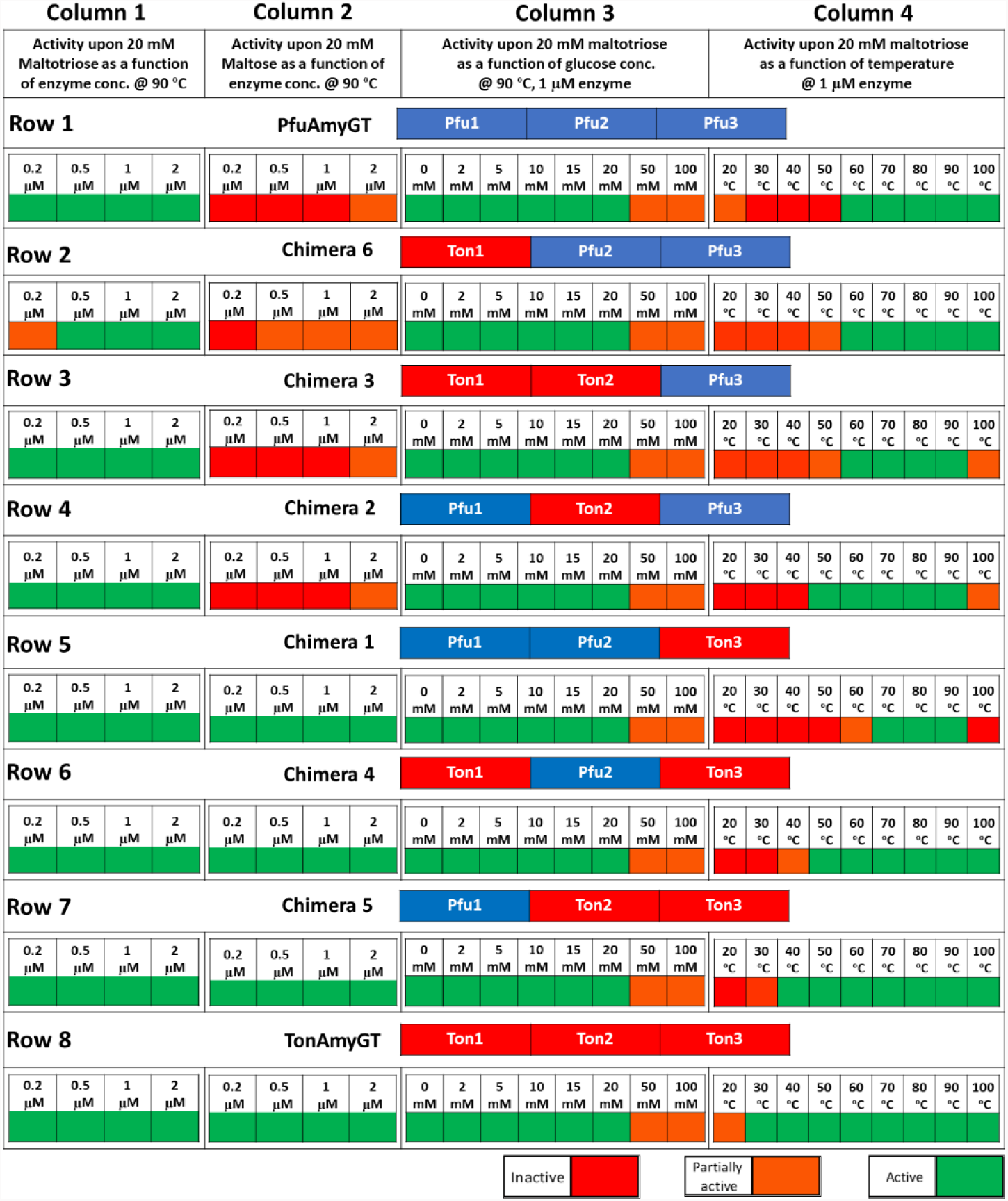
Grid showing the consolidated data of Fig. 6 and 7.

**Fig. 9.**
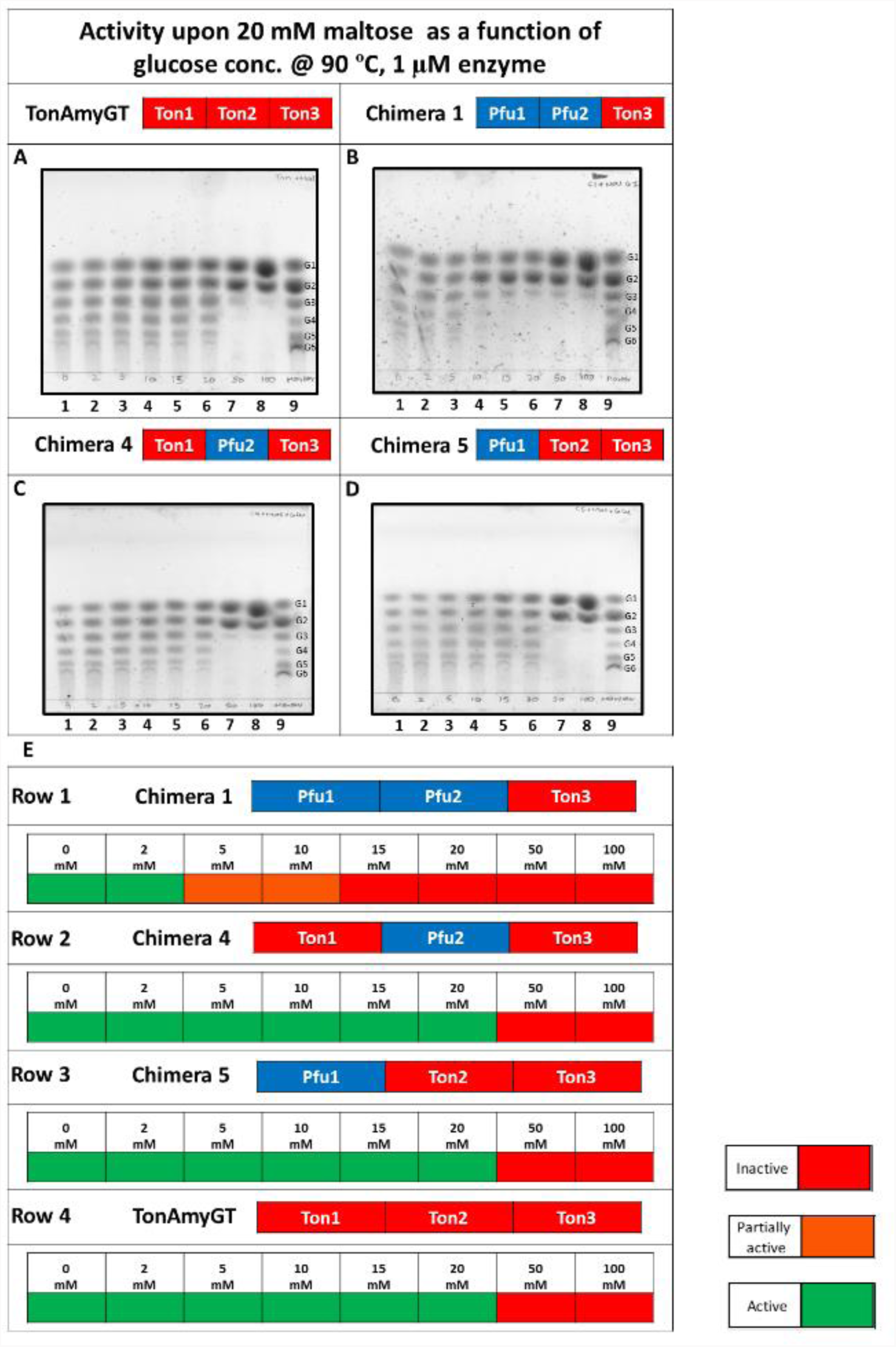
TLC experiments examining the activity profile of TonAmyGT, Chimera 1,4 and 5. **A, B, C And D** TLCs are showing the reaction product formed through 12 h incubations of chimera (1 µM) with maltose (20 mM) at 90 °C as a function of glucose concentration. *Lane 1*: maltose with enzyme and no glucose; *Lane 2*: maltose with 2 mM glucose; *Lane 3*: maltose with 5 mM glucose; *Lane 4*: maltose e with 10 mM glucose; *Lane 5*: maltose with 15 mM glucose; *Lane 6*: maltose with 20 mM glucose; *Lane 7*: maltose with 50 mM glucose; *Lane 8* maltose with 100 mM glucose; ; *Lane 9*: premixed oligosaccharide markers containing 20 mM each of glucose (G1),maltose (G2), maltotriose (G3), maltotetraose (G4), maltopentaose (G5) and maltohexaose (G6). **E.** Grid showing data sourced from Fig. 9A, B, C and D, and consolidated.

#### The preferred smallest donor saccharide of Ton1-Ton2-Ton3 is maltose, whereas that of Pfu1-Pfu2-Pfu3 is maltotriose

The data shown in Supplementary Fig. S6F establishes that Ton1-Ton2-Ton3 is able to act over a wide range of enzyme concentrations (including concentrations as low as 0.2 µM enzyme and as high as 2.0 µM) to transform 20 Mm maltose into a pool of malto-oligosaccharides, over 12 h and at 90 ⁰C, with equilibrium between saccharides of different lengths in this pool being achieved, through dis-proportionating glucanotransfer that lengthens some saccharides while shortening others. This data is summarized in Column 2 in Row 8, in Fig. 8, by boxes coloured green, for reactions involving 0.2, 0.5, 1.0 and 2.0 µM TonAmyGT and 20 mM maltose.

The data shown in Supplementary Fig. S6B establishes that under entirely identical experimental conditions, and using an identical range of enzyme concentrations, Pfu1-Pfu2-Pfu3 is unable to transform 20 mM maltose into the same pool of oligosaccharides. This indicates that Pfu1-Pfu2-Pfu3’s ability to act upon maltose is significantly lower than that of Ton1-Ton2-Ton3 (if indeed Pfu1-Pfu2-Pfu3 is able to use maltose as a substrate at all). This data is summarized in Column 2 in Row 1, in Fig. 8, by boxes that are orange or red for the use of 0.2, 0.5, 1.0 and 2.0 µM Pfu1-Pfu2-Pfu3 with 20 mM maltose.

However, this difference in the catalytic capabilities of the two native enzyme constructs is not observed when maltotriose is given to the enzyme as sole substrate, instead of maltose. With maltotriose, as demonstrated by the TLCs in Supplementary Fig. S6A, for Pfu1-Pfu2-Pfu3, and in Supplementary Fig. S6E, for Ton1-Ton2-Ton3, respectively, both enzymes appear to be capable of transforming maltotriose into a pool of saccharides of different lengths with comparable ease and facility. This data is summarized in Column 1 in Fig. 8, in Rows 1 and 8, respectively, for Pfu1-Pfu2-Pfu3, and Ton1-Ton2-Ton3, using boxes coloured green for the use of 0.2, 0.5, 1.0 and 2.0 µM enzyme and 20 mM maltose. Therefore, we conclude that Ton1-Ton2-Ton3 uses maltose as its smallest donor saccharide whereas Pfu1-Pfu2-Pfu3 uses maltotriose as its smallest donor saccharide, at least over enzyme concentrations ranging from 0.2 to 2.0 µM.

#### Pfu1-Pfu2-Pfu3 can also use maltose as a substrate but only when very high enzyme concentrations are used

Supplementary Fig. S6I shows that when enzyme concentration is raised beyond 2.0 µM, e.g., to 5.0, 10.0 or 20.0 µM, there is detectable activity of Pfu1-Pfu2-Pfu3 seen upon maltose. This indicates that Pfu1-Pfu2-Pfu3’s efficiency of using maltose as a donor substrate is so poor in comparison with that of Ton1-Ton2-Ton3 that detectable levels of conversion of maltose into the pool of saccharides is seen only when enzyme concentration is raised by over one order of magnitude, from 0.2 µM (for Ton1-Ton2-Ton3) to well over 2.0 µM (for Pfu1-Pfu2-Pfu3). This data further corroborates the differences observed between the preferred donor substrate of the two native enzyme constructs, which proves to be useful in interpreting data involving the chimeras, as presented below.

#### All six chimeras display glucanotransferase activity with maltotriose, but to different extents

Figs. 6A, 6E, 6I, 7A, 7E, and 7I, respectively, present raw TLC data for the activities of chimeras 6, 3, 2, 1, 4 and 5, upon maltotriose. Rows 2-7 in Column 1 in Fig. 8 consolidate and summarize this data. We would like to mention here that the odd sequence of presentation of the data for chimeras in Fig. 6-8, i.e., use of the order of 6, 3, 2, 1, 4 and 5 (instead of 1, 2, 3, 4, 5, and 6), was predicated by the manner in which we found it convenient to group chimeras together in Fig. 8. From the summarized data, it is clear that all six chimeras are able to display glucanotransferase activity upon maltotriose (represented by green boxes), with only chimera 6 showing a somewhat compromised ability to do so at 0.1 µM enzyme concentration (represented by an orange box).

#### Chimeras 6, 3, and 2, display compromised glucanotransferase activity with maltose, but chimeras 1, 4 and 5 display normal glucanotransferase activity with maltose

Figs. 6B, 6F, 6J, 7B, 7F, and 7J, respectively, present raw TLC data for the activities of chimeras 6, 3, 2, 1, 4 and 5, upon maltose (which is one glucose shorter in length than maltotriose). Rows 2-7 in Column 2 in Fig. 8 consolidate and summarize this data from which it is clear that chimeras 1, 4, and 5, display normal glucanotransferase activity with maltose (represented by green boxes) whereas chimeras 6, 3, and 2, display grossly compromised levels of activity with maltose (represented by orange and red boxes).

#### Only chimeras incorporating domain 3 from TonAmyGT are able to use maltose as the smallest donor

An analysis of Rows 1 to 4 in column 2 in Fig. 8 shows that there is greatly compromised activity upon maltose in some chimeras, e.g., chimeras 6, 3, and 2, incorporating domain 3 from PfuAmyGT. This is highly significant in the light of the fact that Pfu1-Pfu2-Pfu3 itself also shows greatly compromised activity upon maltose. All four rows in column 2 of Fig. 8 are populated by orange or red boxes, showing that there is little or no activity upon maltose when domain 3 is sourced from Pfu1-Pfu2-Pfu3. In stark contrast, rows 5-8 of column 2 in Fig. 8 are populated by only green boxes, showing that normal glucanotransferase activity is seen in Ton1-Ton2-Ton3 and all chimeras that source their domain 3 from TonAmyGT, e.g., chimeras 1, 4, and 5.

Our summary conclusion from this data is that domain 3 influences the nature and specificity of the smallest donor used by any of these native or chimeric enzyme constructs. Since domain 3, as discussed in the introductory section, is involved (along with domain 2) in creating a secondary/surface substrate-binding site (SBS) on Pfu1-Pfu2-Pfu3 to which maltose was found to be bound, we propose that our findings suggest that the SBS influences substrate specificity. It may be noted that there are other studies involving certain other enzymes in which such SBS sites have been proposed to influence substrate specificity (reference).

#### When using maltose, or maltotriose as donor, all eight constructs are inhibited by high concentrations of glucose, while chimera 1 is inhibited by low concentrations

Raw data is shown in Supplementary Fig. S6C for Pfu1-Pfu2-Pfu3, Fig. 6C (row 2) for Chimera 6, Fig.6G (row 3) for Chimera 3, Fig. 6K (row 4) for Chimera 2, Fig. 7C (row 5) for Chimera 1, Fig. 7G (row 6) for Chimera 4, Fig. 7K (row 7) for Chimera 5, and Supplementary Fig. S6G for Ton1-Ton2-Ton3, respectively, for the action of these constructs upon 20 mM maltotriose. These pieces of raw data are consolidated and summarized in Column 3 of Fig. 8. From this summary, it can be seen that both native constructs and all six chimeric constructs are able to function in the presence of glucose, at concentrations varying from 0 to 20 mM glucose (represented by green boxes), with only some reduction in activity observed at 50 mM and 100 mM glucose (represented by orange boxes).

When everything else remains the same as in the above experiments, and the only difference introduced is that the donor substrate is changed from maltotriose to maltose, the raw data for glucose-based inhibition (for all constructs displaying any ability whatsoever to function upon maltose) is shown in Fig. 9 for Ton1-Ton2-Ton3. The summary of this raw data, shown in Fig. 9E, suggests two things: (i) All four constructs display greater glucose-based inhibition of the activity of enzyme constructs upon maltose (represented by red boxes, seen in four rows of column 3 in Fig. 8) than upon maltotriose (represented by green boxes, seen in all eight rows of column 3 in Fig. 8); (ii) one construct in particular, i.e., Chimera 1, is especially susceptible to glucose-based inhibition of activity upon maltose, with the presence of even 5 mM glucose appearing to compromise activity. Notably, Chimera 1 is the only construct which has a single domain of TonAmyGT out of all four constructs displaying activity with maltose, and this could have something to do with its uniqueness of behaviour in respect of the construct’s inhibition by glucose.

A further perspective on the above observation may be derived from the fact that glucose is closer in size to maltose than it is to maltotriose. Therefore, if glucose competes for binding to an enzyme site with maltose and maltotriose, e.g., at the secondary/surface substrate-binding site (SBS) at the interface of domains 2 and 3, its ability to inhibit action upon maltose could be anticipated to be greater than its ability to inhibit action upon maltotriose. Further, glucose’s inhibiting effect upon the action of enzymes upon maltose could be expected to be manifested to the greatest degree in enzyme constructs possessing some key contribution (e.g., domain 3) from the enzyme (e.g., TonAmyGT) capable of using maltose as a substrate. Also, glucose could be anticipated to maximally affect the enzyme construct characterized by the greatest handicap relating to that specific domain (i.e., domain 3) which plays a key role in the use of maltose as a substrate. Therefore, it appears to us that Chimera 1 shows the greatest inhibiting effect of glucose upon its activity upon maltose, because this is the only chimera amongst all of the maltose-using constructs (i.e., Ton1-Ton2-Ton3, Chimera 1, Chimera 4 and Chimera 5) with the minimum contribution from the native sequence of Ton1-Ton2-Ton3.

#### Pfu1-Pfu2-Pfu3 and Ton1-Ton2-Ton3 display activity over a wide range of temperatures, but some chimeras require higher temperatures to display activity, indicating higher native structural content at higher temperatures

Figs. 6D, 6H, 6L, 7D, 7H, and 7L and Supplementary Figs. S6D, and S6H, respectively, present raw data for the activities of Chimeras 6, 3, 2, 1, 4, 5, Pfu1-Pfu2-Pfu3 and Ton1-Ton2-Ton3 over a wide range of temperatures between 20 and 100 ᵒC. It may be noted that the above figures do not provide us with any indications of differences in the actual catalytic rates of the different enzymes and their chimeras at different temperatures. Rather, the dis-proportionating glucanotransferase reaction produces multiple saccharide species of different lengths in a manner that displays no further change after reaching equilibrium, with this equilibrium being achieved quite rapidly over a time scale of 1 to 2 h, at the enzyme concentrations and maltotriose concentrations used, after which no further change is seen in the relative intensities of the different saccharide bands in the TLC. Therefore, these figures derived from 12 h reactions can be taken to be providing information only about whether the enzyme is functional, and also the extent to which it is functional, i.e., whether it is able to produce all species or only a few species of oligosaccharides, but no information about the catalytic rate(s) that are associated with such functionality.

The raw data is further summarized in column 4 of Fig. 8. The first result that is obvious from this summary is that there are differences in the temperature ranges over which Pfu1-Pfu2-Pfu3, Ton1-Ton2-Ton3, and the six chimeras display activity. All eight enzymes/constructs are fully active at 70, 80 and 90 ᵒC. However, at both lower and higher temperatures, significant differences are seen. Chimera 1 appears to have the narrowest temperature range of function, being completely inactive at temperatures below 70 ᵒC and above 90 ᵒC. Since all domains in the six chimeras are derived from hyperthermophile enzymes, it may be anticipated that high temperatures approaching the growth temperatures of the archaea that naturally produce these enzymes are required to aid the occurrence of folding and assembly into native (and functional) structure, in these chimeras. Depending upon the relative ease of assembly of domains drawn from different parent enzymes, it appears that barring Ton1-Ton2-Ton3which shows full activity at 30 ᵒC and some activity at 20 ᵒC, neither Pfu1-Pfu2-Pfu3, nor any of the chimeras, shows any activity at 30 ᵒC.

Chimera 5, which contains 2 contiguous domains from TonAmyGT (domains 2 and 3), is observed to display full activity at 40 ᵒC. Chimera 4, which contains 2 non-contiguous domains from TonAmyGT (domains 1 and 3), is observed to display partial activity at 40 ᵒC. These two observations involving chimeras with two domains from TonAmyGT indicate a strong inheritance of behaviour linked with temperature-dependence of folding and function from TonAmyGT. All other chimeras and Pfu1-Pfu2-Pfu3 display full activity only at 50 ᵒC and above. Three constructs, in particular, i.e., Chimera 3, Chimera 2 and Chimera 1, appear to have compromised activity at both the higher temperature of 100 ᵒC, and at lower temperatures, e.g., 50 ᵒC and below. Out of these, one chimera in particular, i.e., Chimera 1 appears to be severely compromised in activity at 50 ᵒC and 60 ᵒC, and displays full activity only over a narrow range of temperatures, at 70 ᵒC, 80 ᵒC and 90 ᵒC. Notably, both Pfu1-Pfu2-Pfu3 and Ton1-Ton2-Ton3 and three out of the six chimeras display activity at 100 ᵒC. One obvious correlate of the above data is that lower temperatures of function (e.g., 30 ⁰C or 40 ⁰C) are seen only in chimeras in which domain 3 is sourced from TonAmyGT, whereas higher temperatures of function (e.g., 50 ⁰C or 60 ⁰C) appear to be required for activity to be seen in chimeras in which domain 3 is sourced from PfuAmyGT.

#### All six chimeras display activity upon starch

Supplementary Fig. S7 shows that all six chimeras are capable of utilizing starch as a substrate, similar to the coupled ability to process starch as an amyase as well as maltotriose or maltose as a disproportionating glucanotransferase, seen in full-length PfuAmyGT and TonAmyGT.

#### Insights from subunit-dissociation and unfolding experiments coupled with activity studies on PfuAmyGT-1-2-3

We are also interested in understanding why these enzymes are dimeric in nature. Our primary question in this respect is whether the enzyme is obligatorily homodimeric, i.e., whether there is any activity still present if the enzyme is turned into a monomer. The only way of addressing such a question is to try and dissociate the said homodimer into a monomer; however, without allowing the monomer to undergo any unfolding, to examine whether the monomer is still active. The ionic denaturant, guanidium hydrochloride (Gu-HCl), is useful for addressing such questions, since it functions more as an electrolyte (i.e., like any other salt) than as a chaotropic (disorder-inducing) agent, at low concentrations, but more as a chaotrope than as an electrolyte at higher concentrations. Since the subunit interfaces of proteins are often stabilized by ionic interactions, low concentrations of Gdm.HCl can potentially achieve subunit dissociation without achieving subunit unfolding in some proteins. In such cases, if Gdm.HCl does not interfere with the chemistry of the protein’s function, it become also possible to investigate the presence, or absence of activity, in the dissociated form of the enzyme.

In Supplementary Figs. S8A, B and C, we show that the enzyme construct, PfuAmyGT-1-2-3 is active at concentrations ranging from 0.1 M Gu-HCl to 1 M, 1.2 M, 1.4 M, 1.6 M, 1.8 M, 2 M, 2.5 M and 5 M Gu-HCl, and at all intermediate concentrations of Gu-HCl varying in concentration by increments of 0.1 M Gu-HCl. At 1.0 M Gu-HCl, there is a drastic lowering of activity observed, amounting to a disappearance of activity under the conditions used (i.e., 90 ⁰C, 0.1 µM enzyme, 20 mM maltotriose). Gel filtration elution studies presented in Supplementary Fig. S8D, however, suggest that the enzyme shows a change in elution volume from ∼12 ml (dimer) to 13.6 ml (monomer) at 0.1 M Gu-HCl, indicating that the enzyme remains active as a monomer. When the Gu-HCl concentration is raised to 1.0 M, and the activity is compromised, there seems to be a concomitant change in gel filtration signalling an initiation of unfolding-related perturbations if not outright unfolding, since the elution volume begins to reduce once again beyond this Gu-HCl concentration (suggesting increase, rather than decrease, in hydrodynamic volume). This observation of change in hydrodynamic radii was also corroborated with Dynamic Light Scattering (DLS). In Supplementary Fig. S8E a subtle change in hydrodynamic radii was observed when PfuAmyGT was incubated with and without 100 mM G-HCl. Also, similar observations are made when the enzyme is subjected to Gu-HCl (0.1 M, 1 M, 2 M and 5 M) and incubated at 25 °C overnight as seen from Supplementary Fig. S8F.

## Discussion and Conclusions

Through the exhaustive and comprehensive experimental results presented in the section above, we establish that even if domain 1 hosts a substantial part of the primary catalytic activity seen in GH57 glucanotransferases, a stand-alone form of domain 1 is folded but displays no activity when domains 2 and 3 (which are domains of unknown function) are absent. Further, we show that even when domain 2 is present with domain 1 in a folded construct, there is still no activity observed, although there is activity observed when domain 3 is also present along with domains 1 and 2. This suggests that domain 3, which has been pointed out by us to very likely host a second binding site (SBS) for glucans with catalytic involvement,^20^ and domain 2, which has been pointed out by us to host a second catalytically active aspartate residue,^20^ are both required for activity. The involvement of domain 3 in activity through its hosting of an SBS is fully corroborated by detailed studies of six different chimeras recombining the domains of two different homologous GH57 glucanotransferases (PfuAmyGT and TonAmyGT) which establish that the nature of the observed catalytic activity is substantively determined by the source of domain 3 in the functional chimeric enzyme, with similar preferences of donor substrates for the glucanotransferase reaction being observed in all enzymes that host domain 3 from a particular source, i.e., either PfuAmyGT or TonAmyGT.

## Supporting information

All supplementary tables and figures compiled

## Acknowledgement

We acknowledge funding for this study from the following sources: (1) Centre for Protein Science, Design and Engineering (MHRD-14-0064) from the Ministry of Human Resource Development (MHRD), Government of India, (2) Hyperthermophile Enzyme Hydrolase Research Centre from the Department of Biotechnology (DBT), Government of India, and (3) Intramural funding from the Indian Institute of Science Education and Research (IISER) Mohali. We also acknowledge the significant contributions of Bharathraj Kasilingam (MS thesis student) to some experiments with one of the chimeras (chimera 4).

## Author Credit Statement

AS, PK and PG participated in the conception of the experiments. PK has executed the experiments described in Fig. 3 and Supplementary Fig. S4. AS executed all other experiments. AS contributed towards analysis of data and writing up of the manuscript. PG supervised the design and execution of experiments, analysis of data, and wrote the manuscript along with AS.

