## Supplementary material for "Differences in the activities of domain-swapped chimeras of two homologous GH57 glucanotransferases suggest that a glucan-binding DUF could influence donor substrate specificity": All supplementary tables and figures compiled

**Table ST1**

# A

| | Domain combination | Restriction sites | Vector | Expression host |
|---|---|---|---|---|
| **PfuAmyGT** | Pfu1 Pfu2 Pfu3 | Nde I/ Xho I | pET23a | BL21 (DE3) pLySs star |
| **TonAmyGT** | Ton1 Ton2 Ton3 | Nhe I/Xho I | pET23a | Rosetta (DE3) |
| **Chimera 1** | Pfu1 Pfu2 Ton3 | Nhe I/Xho I | pET23a | Rosetta (DE3) |
| **Chimera 2** | Pfu1 Ton2 Pfu3 | Nde I/Xho I | pET23a | Rosetta (DE3) |
| **Chimera 3** | Ton1 Ton2 Pfu3 | Nhe I/Xho I | pET23a | BL21 (DE3) pLySs star |
| **Chimera 4** | Ton1 Pfu2 Ton3 | Nhe I/Xho I | pET23a | Rosetta (DE3) |
| **Chimera 5** | Pfu1 Ton2 Ton3 | Nhe I/Xho I | pET23a | Rosetta (DE3) |
| **Chimera 6** | Ton1 Pfu2 Pfu3 | Nhe I/Xho I | pET23a | Rosetta (DE3) |

# B

| | Domain combination | Molecular weight of protein () | pI | Abs 0.1% (=1 g/l) |
|---|---|---|---|---|
| **PfuAmyGT** | Pfu1 Pfu2 Pfu3 | 78.1 | 5.58 | 2.049 |
| **TonAmyGT** | Ton1 Ton2 Ton3 | 76.6 | 5.46 | 1.831 |
| **Chimera 1** | Pfu1 Pfu2 Ton3 | 77.9 | 5.44 | 2.069 |
| **Chimera 2** | Pfu1 Ton2 Pfu3 | 77.8 | 5.48 | 2.02 |
| **Chimera 3** | Ton1 Ton2 Pfu3 | 76.5 | 5.63 | 1.81 |
| **Chimera 4** | Ton1 Pfu2 Ton3 | 76.6 | 5.49 | 1.86 |
| **Chimera 5** | Pfu1 Ton2 Ton3 | 77.8 | 5.33 | 2.03 |
| **Chimera 6** | Ton1 Pfu2 Pfu3 | 76.6 | 5.79 | 1.85 |

**Table ST1: ST1A** shows information about the domain combinations, restriction sites in which the genes are cloned, the vector in which the plasmid is expressed and the expression host cells for PfuAmyGT, TonAmyGT and the six chimeras. **ST1B** shows information about the domain combinations, molecular weight of the proteins, pI of the proteins and the absorbance value at which the protein is 1 mg/ml for PfuAmyGT, TonAmyGT and the six chimeras.

**Table ST2**

| | |
|---|---|
| Pfu F (Nde) | 5'-AATAATCATATGATCAATGGTTGGACC-3' |
| Pfu F (Nhe) | 5'-ATATATGCTAGCATGATCAATGGTTGGACCGAAGTTGGTGAC-3' |
| Pfu R | 5'-AATAATCTCGAGACCAGACGCTTC-3' |
| Ton F | 5'-ATATATGCTAGCATGGTCAATTTCATCTTTGG-3' |
| Ton R | 5'-ATATATGCGGCCGCCAACTCCTTAAAGCTGAGCC-3' |
| P1T2 F | 5'-GTATCTTCGAAAAATACCGCGTCTTC-3' |
| P1T2 R | 5'-AAGACGCGGTATTTTTCGAAGATAC-3' |
| P2T3 F | 5'-AAAGCGAACTCTTACGTTTTCACTGGAAGTTTCGTCAG-3' |
| P2T3 R | 5'-CTGACGAAACTTCCAGTGAAAACGTAAGAGTTCGCTTTG-3' |
| T1P2 F | 5'-CAGTTCGAGAAGTACCGTGTTTTCGTTCGTGG-3' |
| T1P2 R | 5'-CACGAACGAAAACACGGTACTTCTCGAACTGGC-3' |
| T2P3 F | 5'-AAGGCGAACAGCTACGTTTCTCTGGGTAAAGTTATC-3' |
| T2P3 R | 5'-GATAACTTTACCCAGAGAAACGTAGCTGTTCGCC-3' |

**Table ST2** shows the list of primers for achieving PfuAmyGT, TonAmyGT and the six chimeras.

**Table ST3**

| | Domain 1 | Domain 2 | Domain 3 | Total |
|---|---|---|---|---|
| TLGT | 7 | 3 | 5 | 15 |
| PfuAmyGT | 12 | 3 | 3 | 18 |
| TonAmyGT | 9 | 3 | 4 | 16 |

**Table ST3** lists the number of tryptophans in each domain of PfuAmyGT, TonAmyGT and TliAmyGT (TLGT).

**Fig. S1**

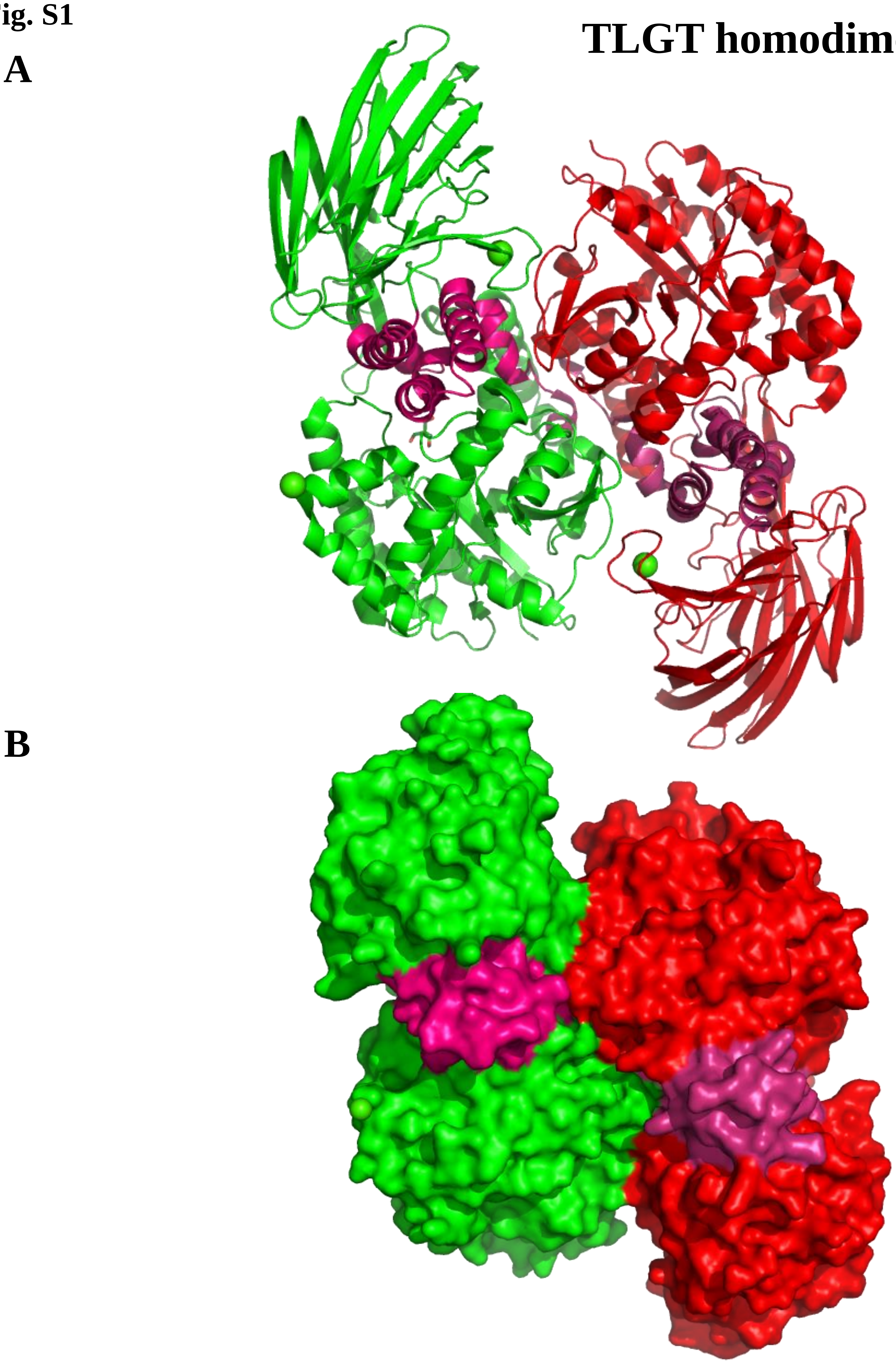

**Fig. S1**. Figure shows the dimeric structure of TLGT. **A and B** shows the ribbon and whole surface representation of TLGT. Red colored subunit is subunit 1 and green colored is subunit 2. Magenta and pink colored domains are domain 2 from subunit 1 and 2 respectively.

Fig. S2

A

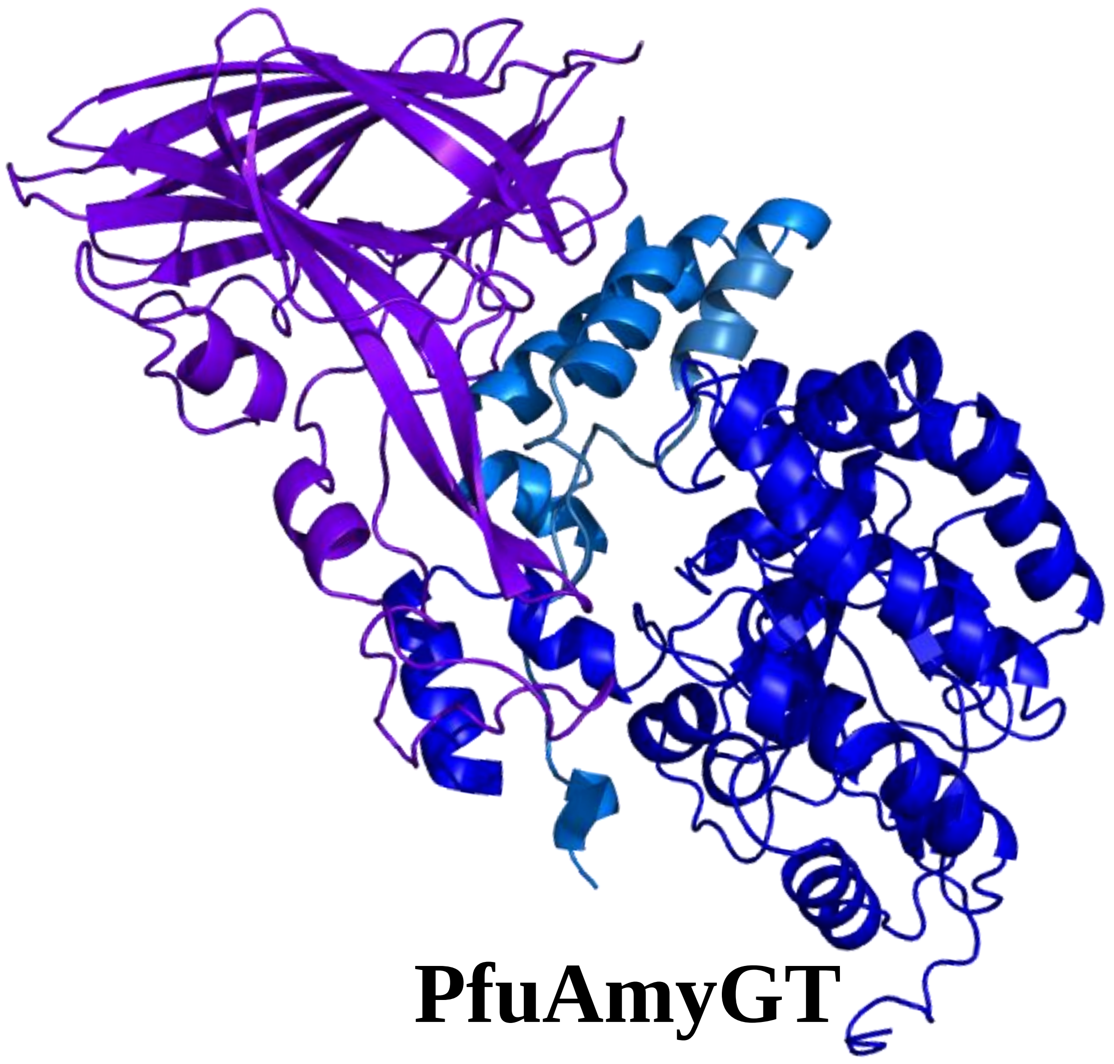

PfuAmyGT

| Domains | Domain color | Domain boundaries |
|---|---|---|
| Domain 1 | | 1-292 |
| Domain 2 | | 312-385 |
| Domain 3 | | 393-658 |

B

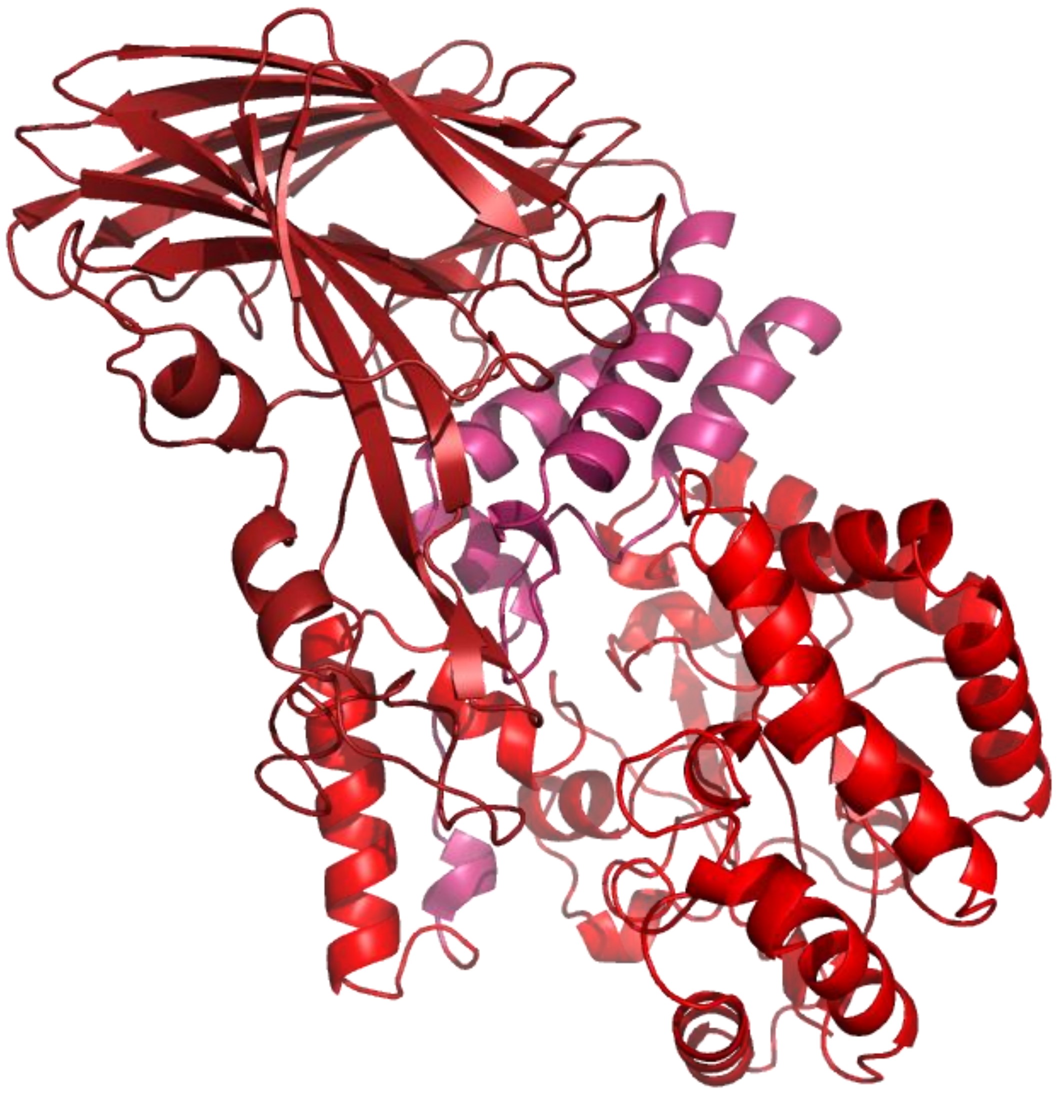

TonAmyGT

| Domains | Domain color | Domain boundaries |
|---|---|---|
| Domain 1 | | 1-282 |
| Domain 2 | | 302-375 |
| Domain 3 | | 383-652 |

C

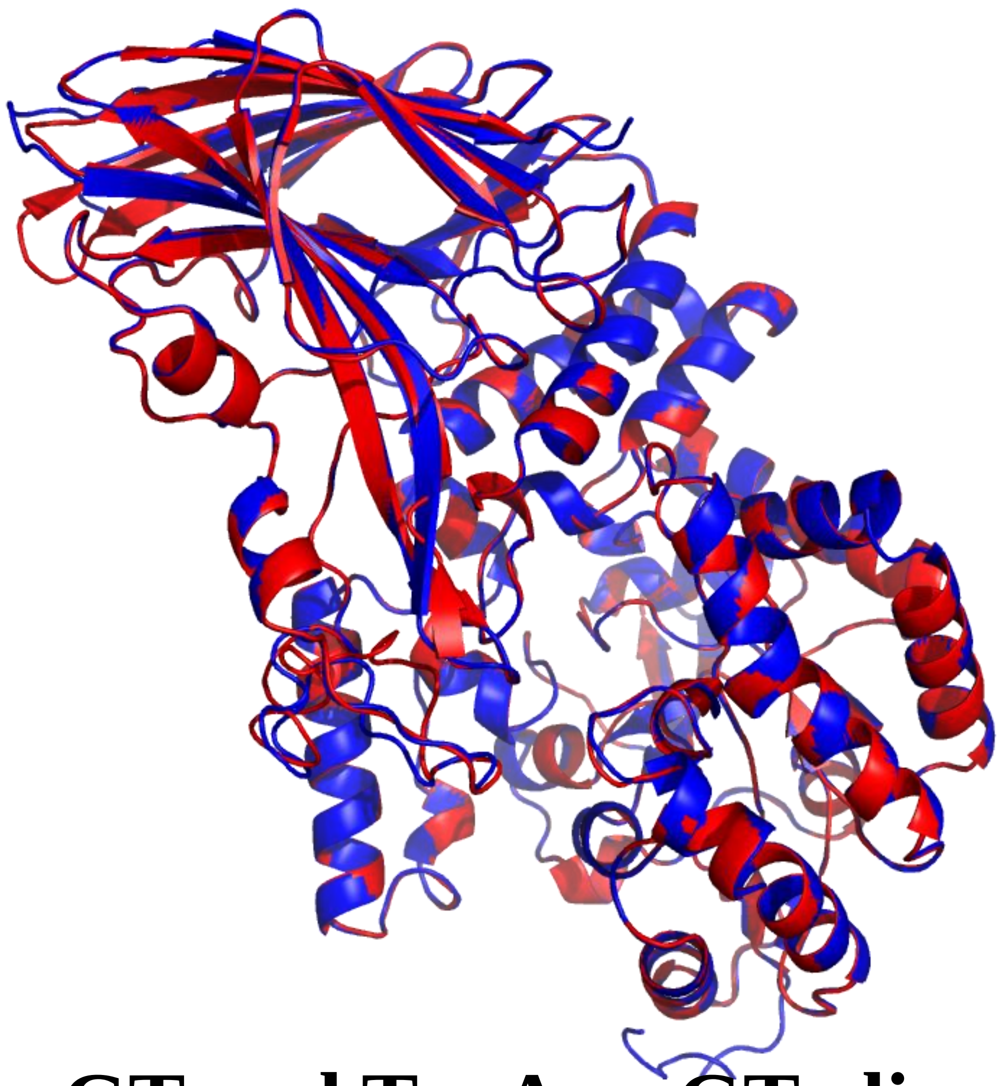

PfuAmyGT and TonAmyGT aligned

| | Percent identity between domains of PfuAmyGT and TonAmyGT |
|---|---|
| Domain 1 | 82.67 % |
| Domain 2 | 86.42 % |
| Domain 3 | 50.93 % |

**Fig. S2**: **Modelled structure of PfuAmyGT and TonAmyGT (through i-Tasser) A** shows the modelled structure of PfuAmyGT with three domains of different color and their respective boundaries (presented in the table bedside the structure). **B** shows the modelled structure of TonAmyGT with three domains of different color and their respective boundaries (presented in the table bedside the structure). **C s**hows the superimposed structure of PfuAmyGT and TonAmyGT (RMSD- 0.2) and the table is showing the Percent identity between domains of PfuAmyGT and TonAmyGT.

**Fig. S3**

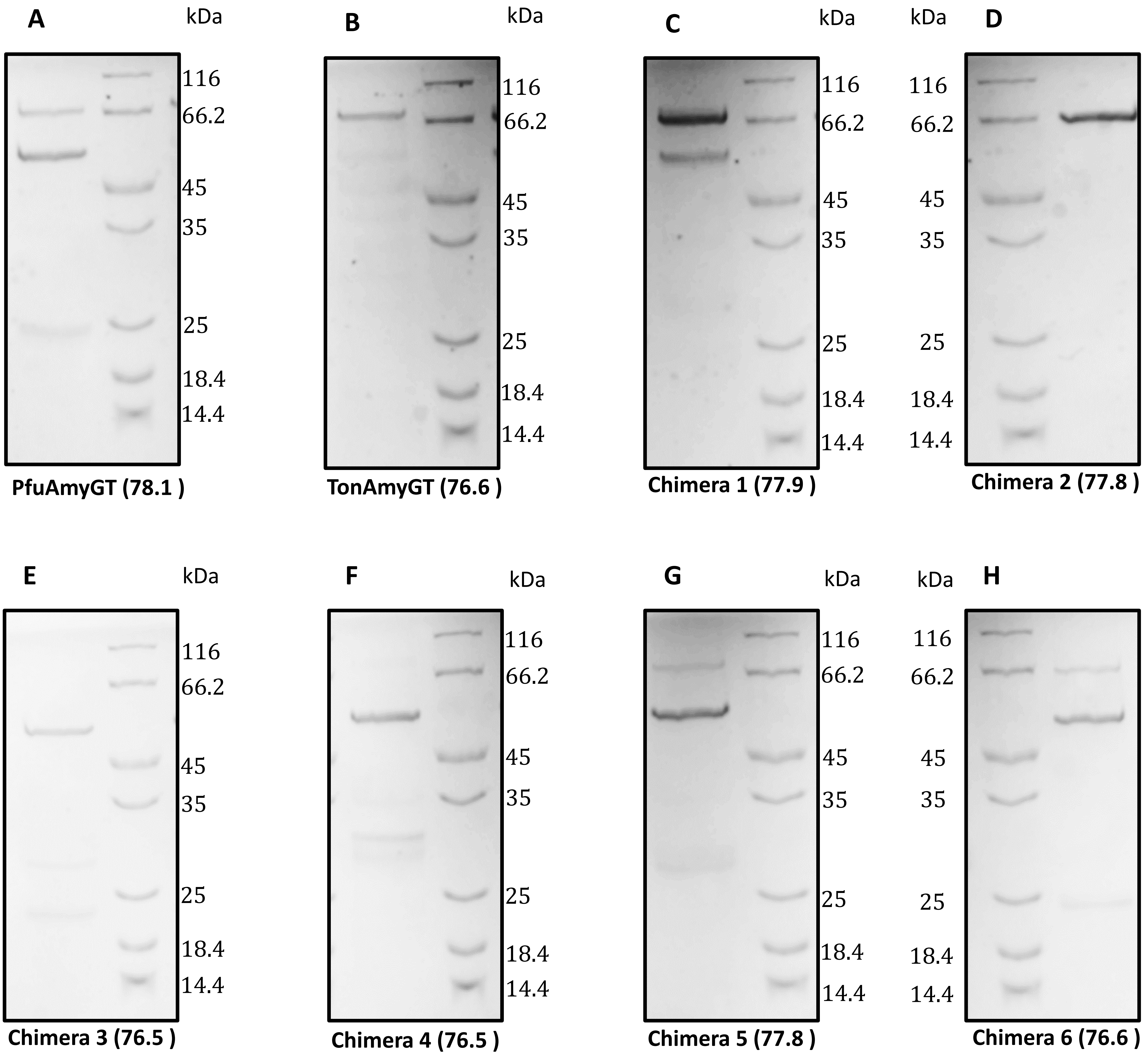

**Fig. S3.** The figure shows the expression gels of the chimeras after second step of purification through Size Exclusion Chromatography.

**Fig. S4**

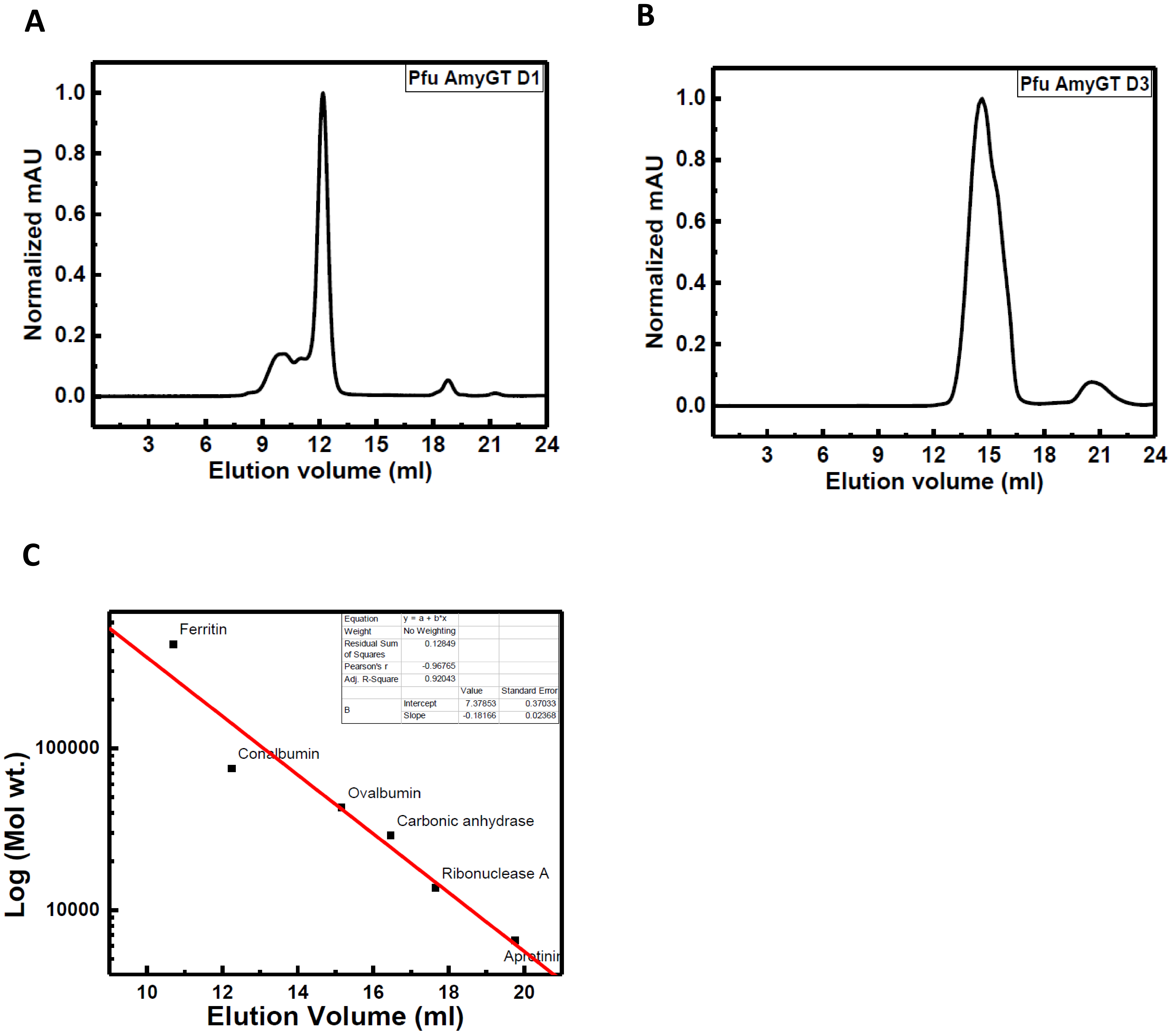

**Fig. S4: Gel filtration chromatogram of two domains: A and B** shows the gel filtration chromatogram of Domain 1 and Domain 3 respectively. **C** shows the calibration graph for Superdex-200 Increase 10/300 GL column with various molecular weight standard proteins.

**Fig. S5**

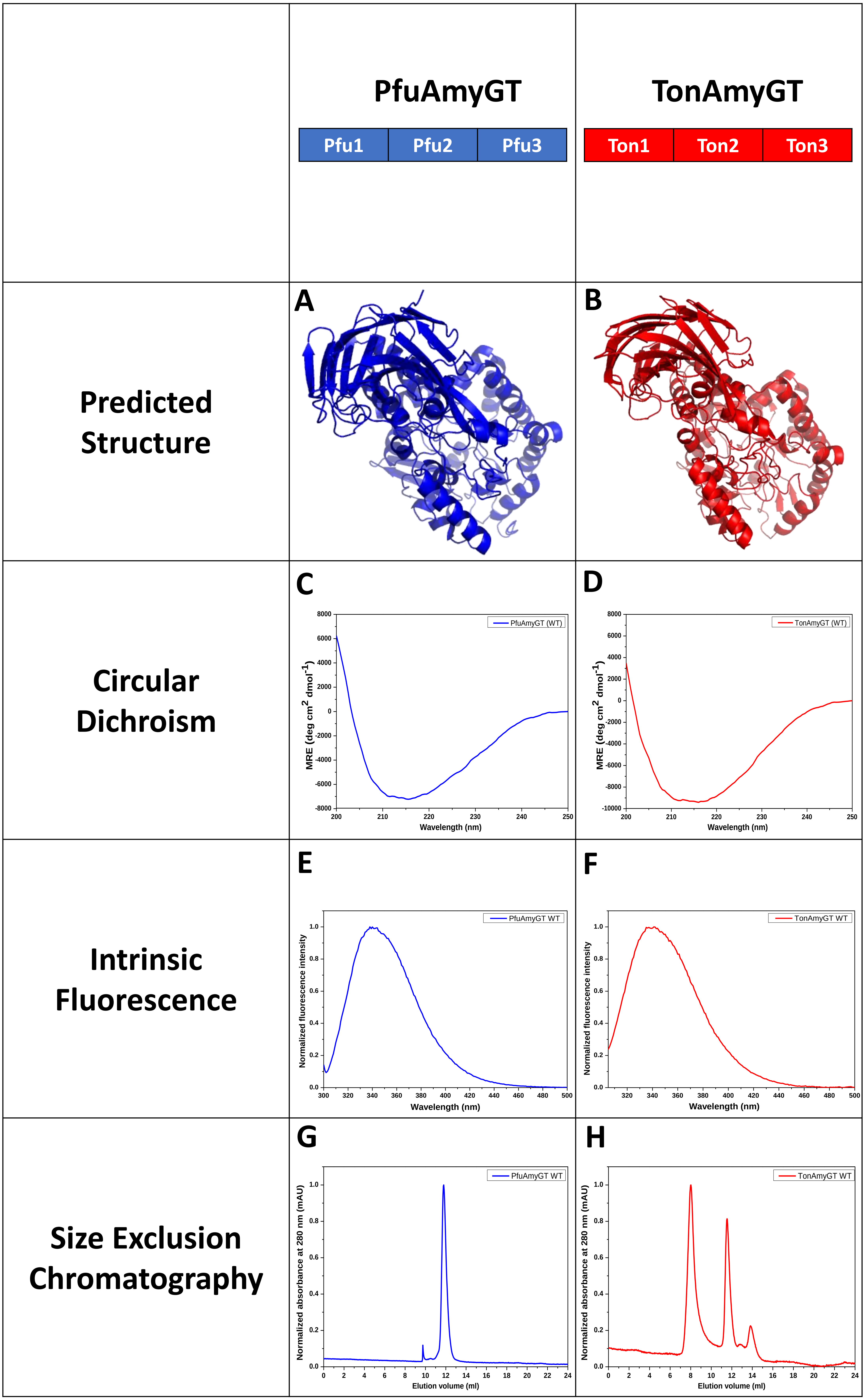
PfuAmyGT
Pfu1
Pfu2
Pfu3
TonAmyGT
Ton1
Ton2
Ton3
Predicted Structure
A
B
Circular Dichroism
C
D
MRE (deg cm2 dmol-1)
Wavelength (nm)
PfuAmyGT (WT)
TonAmyGT (WT)
Intrinsic Fluorescence
E
F
Normalized fluorescence intensity
Wavelength (nm)
PfuAmyGT WT
TonAmyGT WT
Size Exclusion Chromatography
G
H
Normalized absorbance at 280 nm (mAU)
Elution volume (ml)
PfuAmyGT WT
TonAmyGT WT

**Fig. S5. Comparative analyses of predicted structure, spectroscopic and activity for PfuAmyGT and TonAmyGT. A and B** shows modelled structure of PfuAmyGT and TonAmyGT**, C and D** Circular Dichroism (CD) spectra of PfuAmyGT and TonAmyGT**, E and F** Normalized fluorescence emission spectra of PfuAmyGT and TonAmyGT**, G and H** Normalized gel filtration chromatogram of the PfuAmyGT and TonAmyGT.

### Fig. S6

| Activity upon 20 mM Maltotriose as a function of enzyme conc. @ 90 °C | Activity upon 20 mM Maltose as a function of enzyme conc. @ 90 °C | Activity upon 20 mM maltotriose as a function of glucose conc. @ 90 °C, 1 μM enzyme | Activity upon 20 mM maltotriose as a function of temperature @ 1 μM enzyme |
|---|---|---|---|

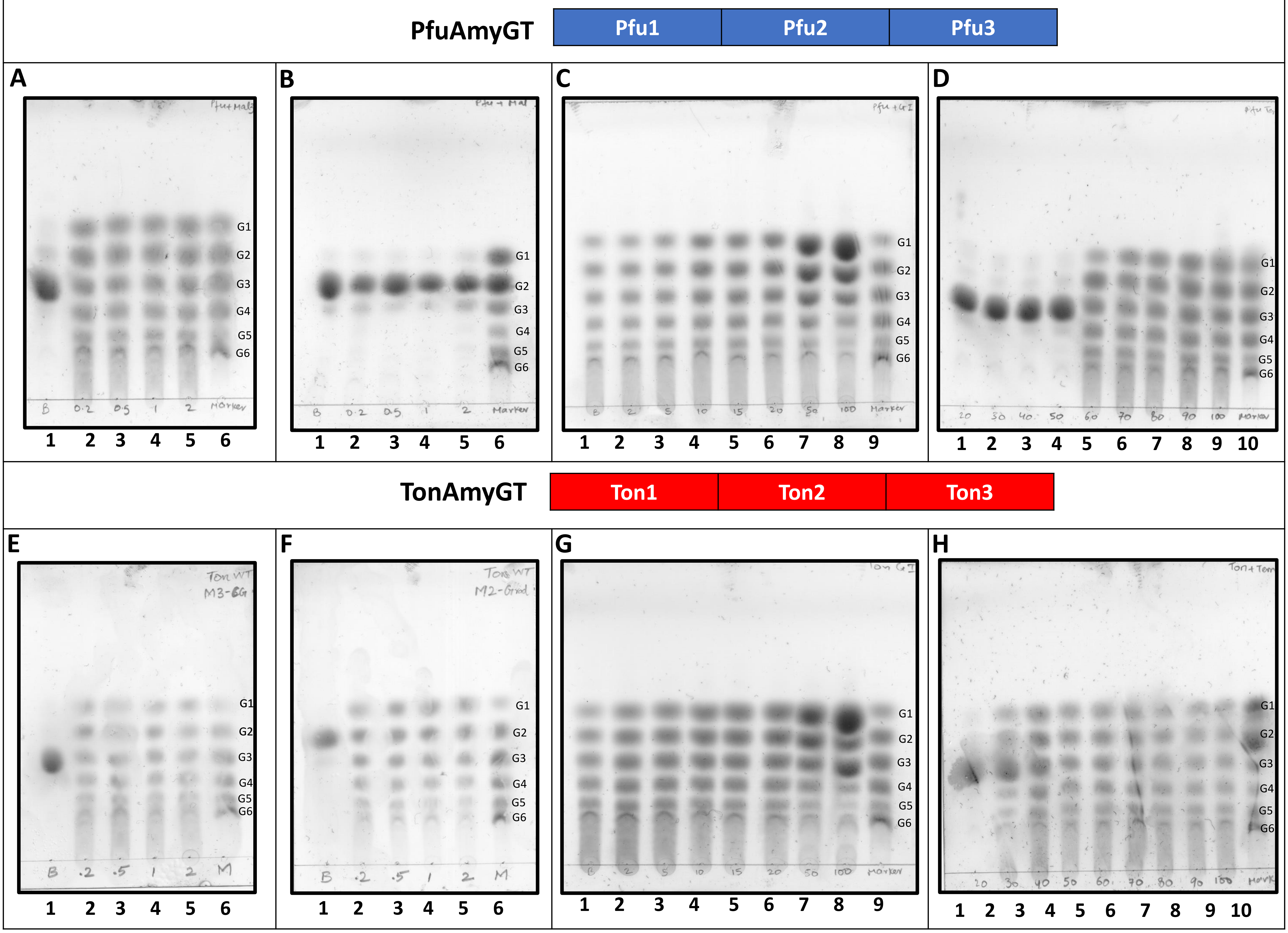

I

Activity upon 20 mM Maltose as a function of enzyme conc. @ 90 °C

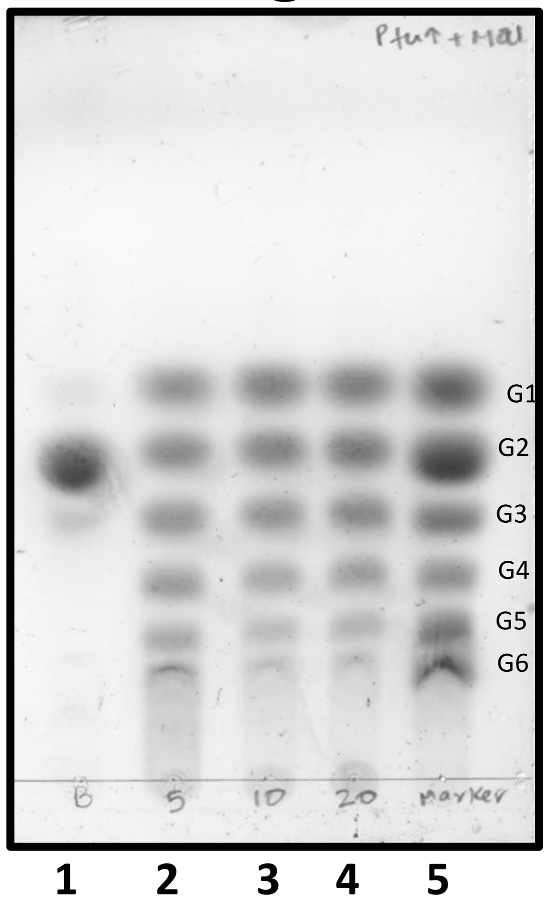

**Fig. 6. TLC experiments examining the activity profile of Chimera 1,5 and 6. A and E** TLCs are showing the reaction products formed through 12 h incubations of PfuAmyGT and TonAmyGT respectively, with maltotriose(20 mM) at 90 °C. *Lane 1*: maltotriose with no enzyme; *Lane 2*: maltotriose with 0.2 μM enzyme; *Lane3*: maltotriose with 0.5 μM enzyme; *Lane 4*: maltotriose with 1 μM enzyme; *Lane 5*: maltotriose with 2 μM enzyme; *Lane 6*: premixed oligosaccharide markers containing 20 mM each of glucose (G1),maltose (G2), maltotriose (G3), maltotetraose (G4), maltopentaose (G5) and maltohexaose (G6). **B and F** TLCs are showing the reaction products formed through 12 h incubations of PfuAmyGT and TonAmyGT respectively, with maltose (20 mM) at 90 °C. *Lane 1*: maltose with no enzyme; *Lane 2*: maltose with 0.2 μM enzyme; *Lane3*: maltose with 0.5 μM enzyme; *Lane 4*: maltose with 1 μM enzyme; *Lane 5*: maltose with 2 μM enzyme; *Lane 6*: premixed oligosaccharide markers containing 20 mM each of glucose (G1),maltose (G2), maltotriose (G3), maltotetraose (G4), maltopentaose (G5) and maltohexaose (G6). **C and G** TLCs are showing the reaction product formed through 12 h incubations of PfuAmyGT and TonAmyGT respectively (1 μM) with maltotriose (20 mM) at 90 °C as a function of glucose concentration. *Lane 1*: maltotriose with enzyme and no glucose; *Lane 2*: maltotriose with 2 mM glucose; *Lane 3*: maltotriose with 5 mM glucose; *Lane 4*: maltotriose with 10 mM glucose; *Lane 5*: maltotriose with 15 mM glucose; *Lane 6*: maltotriose with 20 mM glucose; *Lane 7*: maltotriose with 50 mM glucose; *Lane 8* maltotriose with 100 mM glucose; ; *Lane 9*: premixed oligosaccharide markers containing 20 mM each of glucose (G1),maltose (G2), maltotriose (G3), maltotetraose (G4), maltopentaose (G5) and maltohexaose (G6). **D and H** TLCs are showing the reaction product formed through 12 h incubations of PfuAmyGT and TonAmyGT respectively (1 µM) with maltotriose (20 mM) as a function of temperature. *Lane 1*: maltotriose with enzyme at 20 ºC; *Lane 2*: maltotriose with enzyme at 30 ºC; *Lane 3*: maltotriose with enzyme at 40 ºC; *Lane 4*: maltotriose with enzyme at 50 ºC; *Lane 5*: maltotriose with enzyme at 60 ºC; *Lane 6*: maltotriose with enzyme at 80 ºC; *Lane 7*: maltotriose with enzyme at 90 ºC; *Lane 8*: maltotriose with enzyme at 100 ºC; *Lane 9* maltotriose with 100 mM glucose; *Lane 10*: premixed oligosaccharide markers containing 20 mM each of glucose (G1),maltose (G2), maltotriose (G3), maltotetraose (G4), maltopentaose (G5) and maltohexaose (G6). **I** TLC shows the reaction products formed through 12 h incubations of PfuAmyGT with maltose (20 mM) at 90 °C. *Lane 1*: maltotriose with no enzyme; *Lane 2*: maltose with 5 μM enzyme; *Lane3*: maltose with 10 μM enzyme; *Lane 4*: maltotriose with 20 μM enzyme; *Lane 5*: premixed oligosaccharide markers containing 20 mM each of glucose (G1), maltose (G2), maltotriose (G3), maltotetraose (G4), maltopentaose (G5) and maltohexaose (G6).

**Fig. S7**

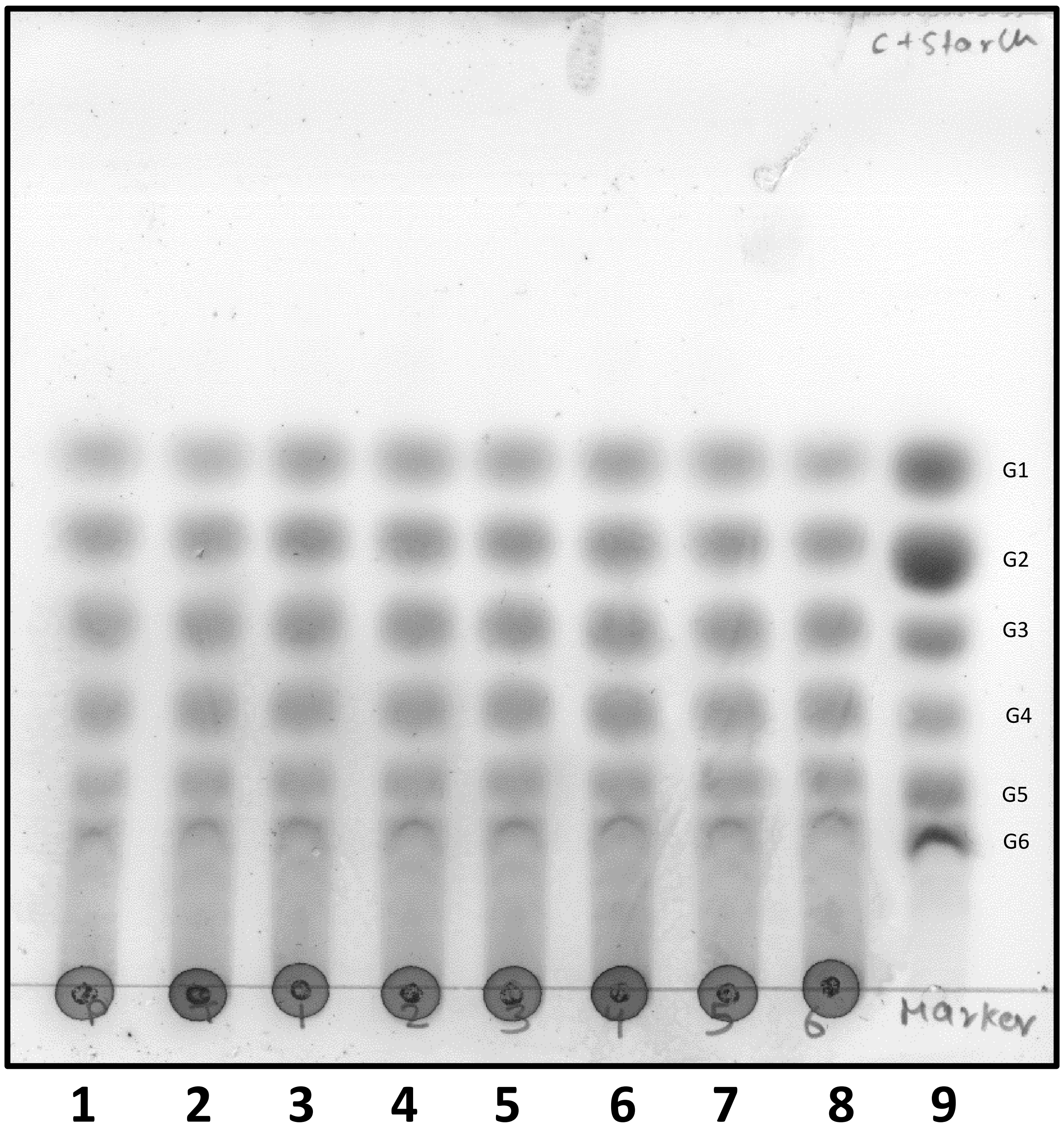

**Fig. S7.** TLCs are showing the reaction products formed through 12 h incubations of PfuAmyGT TonAmyGT (1 µM) and all six chimeras (1 µM each) with 1% starch at 90 °C. *Lane 1*: PfuAmyGT (1 µM) and starch ; *Lane 2*: TonAmyGT (1 µM) and starch; *Lane3*: Chimera 1 (1 µM) and starch ; *Lane 4*: Chimera 2 (1 µM) and starch; *Lane 5*: Chimera 3 (1 µM) and starch; *Lane 6*: Chimera 4 (1 µM) *Lane 7*: Chimera 5 (1 µM) *Lane 8*: Chimera 6 (1 µM); *Lane 9*: premixed oligosaccharide markers containing 20 mM each of glucose (G1), maltose (G2), maltotriose (G3), maltotetraose (G4), maltopentaose (G5) and maltohexaose (G6).

**Fig. S8**

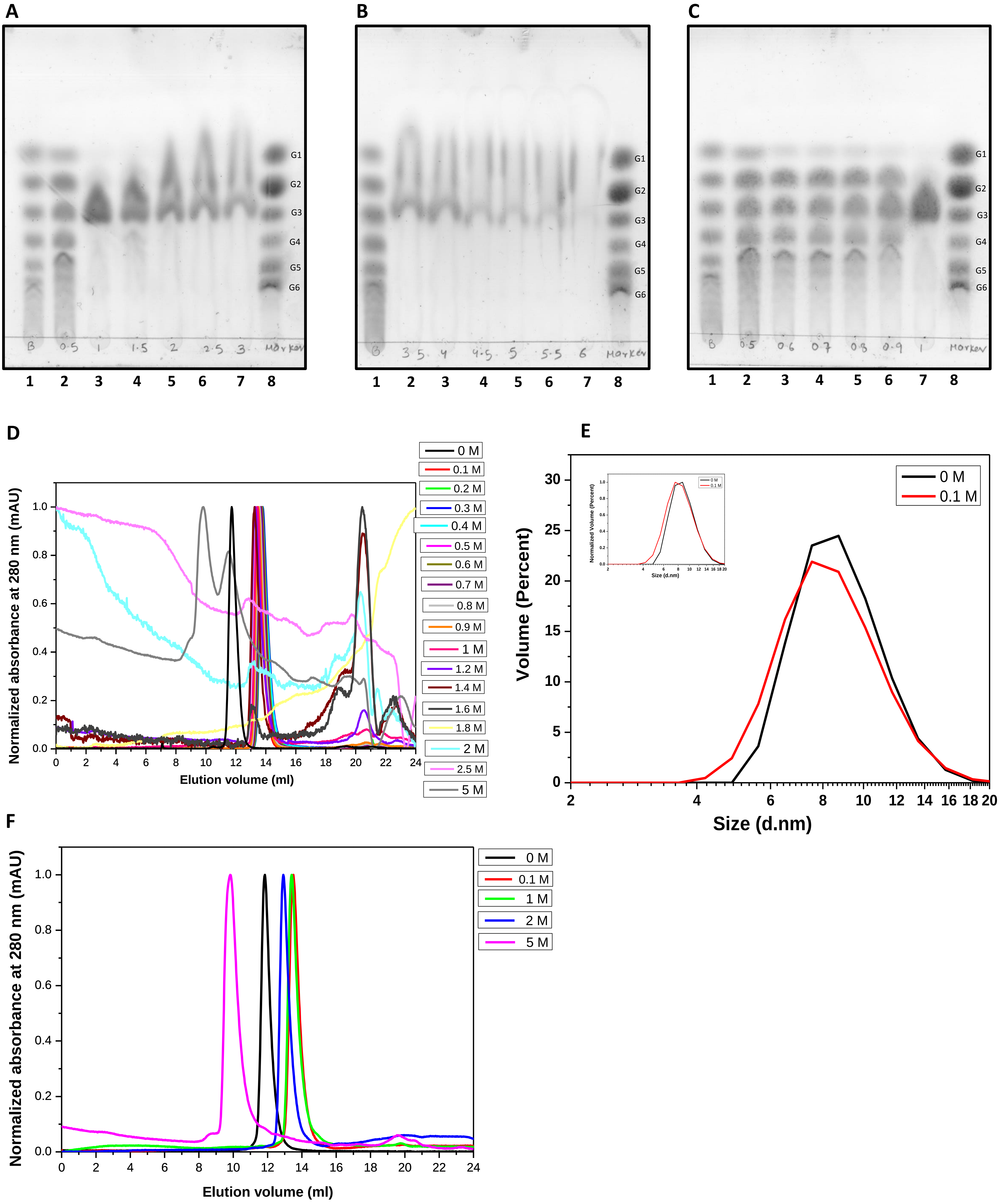
A
B 0.5 1 1.5 2 2.5 3 Marker
G1
G2
G3
G4
G5
G6
1 2 3 4 5 6 7 8
B
B 3.5 4 4.5 5 5.5 6 Marker
G1
G2
G3
G4
G5
G6
1 2 3 4 5 6 7 8
C
B 0.5 0.6 0.7 0.8 0.9 1 Marker
G1
G2
G3
G4
G5
G6
1 2 3 4 5 6 7 8
D
Normalized absorbance at 280 nm (mAU)
Elution volume (ml)
0 M
0.1 M
0.2 M
0.3 M
0.4 M
0.5 M
0.6 M
0.7 M
0.8 M
0.9 M
1 M
1.2 M
1.4 M
1.6 M
1.8 M
2 M
2.5 M
5 M
E
Volume (Percent)
Size (d.nm)
0 M
0.1 M
Normalized Volume (Percent)
F
Normalized absorbance at 280 nm (mAU)
Elution volume (ml)
0 M
0.1 M
1 M
2 M
5 M

**Fig. S8. A, B and C,** TLCs shows the activity of PfuAmyGT in presence of different concentrations of guanidium hydrochloride (Gu-HCl) at 90 °C for 12 h. **D,** shows the SEC chromatogram of PfuAmyGT incubated with Gu-HCl at 90 °C for 12 h. **E**, shows the volume percentage of PfuAmyGT in presence and absence of Gu-HCl, through DLS. **F,** TLCs shows the activity of PfuAmyGT in presence of different concentrations of guanidium hydrochloride (Gu-HCl) incubated at 25 °C for 12 h.
